# A cryptic BRCA2 repeated motif binds to HSF2BP oligomers with no impact on meiotic recombination

**DOI:** 10.1101/2020.12.29.424679

**Authors:** Rania Ghouil, Simona Miron, Lieke Koornneef, Jasper Veerman, Maarten W. Paul, Marie-Hélène Le Du, Esther Sleddens-Linkels, Sari E. van Rossum-Fikkert, Yvette van Loon, Natalia Felipe-Medina, Alberto M. Pendas, Alex Maas, Jeroen Essers, Pierre Legrand, Willy M. Baarends, Roland Kanaar, Sophie Zinn-Justin, Alex N. Zelensky

## Abstract

BRCA2 plays a prominent role in meiotic homologous recombination (HR). Loss of BRCA2 or several of its meiotic partners causes fertility defects. One of these partners, HSF2BP, was recently discovered as expressed physiologically in germline and ectopically produced in cancer cells. It has an N-terminal coiled coil motif involved in direct binding to the protein BRME1, and both HSF2BP and BRME1 are essential for meiotic HR during spermatogenesis. It also interacts through its C-terminal Armadillo (ARM) domain with a conserved region of BRCA2 of unknown function. We analyzed the structural properties and functional consequences of the BRCA2-HSF2BP interaction and tested the emerging model of its involvement in meiosis. We solved the crystal structure of the complex between the BRCA2 fragment that is disordered in solution and the HSF2BP dimeric ARM domain. This revealed two previously unrecognized BRCA2 repeats that each interact with one ARM monomer from two different dimers. BRCA2 binding triggers ARM tetramerization, resulting in a complex containing two BRCA2 fragments connecting two ARM dimers. The 3D structures of the BRCA2 repeats are superimposable, revealing conserved contacts between the BRCA2 residues defining the repeats and the HSF2BP residues lining the groove of the ARM. This large interface is responsible for the nanomolar affinity of the interaction, significantly stronger than any other measured interaction involving BRCA2. Deleting exon 12 from *Brca2*, encoding the first repeat, disrupted BRCA2 binding to HSF2BP *in vitro* and in cells. However, *Brca2^Δ12/Δ12^* mice with the same deletion were fertile and did not show any meiotic defects, contrary to the prediction from the model positing that HSF2BP acts as a meiotic localizer of BRCA2. We conclude that the high-affinity interaction between BRCA2 and HSF2BP and the resulting HSF2BP oligomerization are not required for RAD51 and DMC1 recombinase localization to meiotic double strand breaks and for productive meiotic HR.

## Introduction

Homologous recombination (HR) is involved in many aspects of eukaryotic DNA metabolism and is indispensable in two contexts: resolving replication problems and in meiosis. Homology search and strand exchange, the key events in HR, are performed by a nucleoprotein filament formed by the strand exchange protein RAD51 assembled onto the 3’ single stranded (ss) DNA overhang. In somatic animal cells, RAD51 loading onto ssDNA depends on BRCA2, which has multiple RAD51 binding sites and is required for focal accumulation of RAD51 at the sites of damage. Biochemical experiments suggest that BRCA2 acts as an HR mediator, displacing RPA, the protein that protects ssDNA by strongly binding to it, and forming functional RAD51 filament in its place (reviewed in Zelensky et al., 2014). *In vitro*, BRCA2 can perform this function autonomously, but in cells it depends on its “partner and localizer” PALB2 (Xia et al., 2006).

Although BRCA2 has been mostly studied in the context of HR in somatic cells, it arguably has a more prominent role in meiotic HR, as across a broad range of species, fertility defects are the most common consequence of BRCA2 loss (Connor et al., 1997; Klovstad et al., 2008; Ko et al., 2008; Martin et al., 2005; Miao et al., 2019; Sharan et al., 2004; Shive et al., 2010; Siaud et al., 2004; Weinberg-Shukron et al., 2018). In meiosis, HR functions to diversify as well as to preserve genetic information. This role is achieved by extending the core HR machinery (RAD51, BRCA2, PALB2) with a set of meiosis-specific proteins, such as the DMC1 recombinase and the ssDNA-binding proteins MEIOB and SPATA22. BRCA2 binds DMC1 via the RAD51-binding BRC repeats encoded by *BRCA2* exon 11 (Dray et al., 2006; Martinez et al., 2016; Thorslund et al., 2007) and a DMC1-specific site encoded by exon 14 (Thorslund et al., 2007).

We identified HSF2BP as another BRCA2-binding protein, endogenously expressed in meiotic cells, and ectopically produced in cancer cells (Brandsma et al., 2019, 2016; Sato et al., 2020). HSF2BP is required for meiotic HR during spermatogenesis, but in somatic cells, it inhibits HR during DNA inter-strand crosslink repair by triggering BRCA2 degradation. In addition to BRCA2, HSF2BP has been reported to interact with transcription factors HSF2 (Yoshima et al., 1998) and BNC1 (Wu et al., 2013), both required for normal fertility in mice (Kallio et al., 2002; Zhang et al., 2012). More recently, five groups independently reported that HSF2BP interacts with an uncharacterized protein named C19orf57, 4930432K21Rik, <u>BRME1</u>, MEIOKE21 or MAMERR. The two proteins co-localize in meiocytes, loss of BRME1 closely phenocopies loss of HSF2BP, and the two proteins can affect each other’s stability (Felipe-Medina et al., 2020; Li et al., 2020; Shang et al., 2020; Takemoto et al., 2020; Zhang et al., 2020). The model put forward to explain the meiotic defects in knock out mice for either *Hsf2bp* or *Brme1* follows the PALB2 paradigm: HSF2BP and BRME1 are proposed to act as “meiotic localizers” for BRCA2 — and for each other (Zhang et al., 2020, 2019). However, how HSF2BP, BRME1 and PALB2 contribute to BRCA2 localization in meiotic HR remains to be established. Also, in contrast to complete dependence of BRCA2 on PALB2 in somatic HR, the loss of the two meiotic localizers (HSF2BP and BRME1) causes a milder meiotic phenotype than *Brca2* deficiency in mice. *Hsf2bp* and even more so *Brme1* knockout mouse models show pronounced sexual dimorphism: female meiotic defects are either weak (Felipe-Medina et al., 2020; Zhang et al., 2019) or not detected (Brandsma et al., 2019; Li et al., 2020; Shang et al., 2020; Takemoto et al., 2020; Zhang et al., 2020), while *Brca2* deficiency affects both sexes (Sharan et al., 2004).

HSF2BP contains an N-terminal α-helical oligomerisation domain and a C-terminal domain predicted to adopt an Armadillo (ARM) fold (Brandsma et al., 2019, 2016; Zhang et al., 2020). We mapped the HSF2BP-BRCA2 interaction to the Armadillo domain of HSF2BP, and a 68 amino acid (aa) region of BRCA2 mostly encoded by exons 12 and 13, which is predicted to be disordered (Brandsma et al., 2019). In fact, BRCA2 is a protein of 3,418 aa that possesses a unique globular domain of 700 aa binding to ssDNA and the acidic protein DSS1 (Yang et al., 2002). The high disorder propensity of BRCA2 is proposed to ensure its structural plasticity and its ability to orchestrate complex molecular transactions while balancing multiple interactions (Sánchez et al., 2017; Sidhu et al., 2020). However, interactions involving intrinsically disordered regions are difficult to predict and characterize structurally. Out of more than a dozen mapped protein interaction regions within the disordered part of BRCA2 (Martinez et al., 2015; Prakash et al., 2015), only four were crystallized when bound to their folded partner, either Rad51 (Pellegrini et al., 2002), PALB2 (Oliver et al., 2009) or PLK1 (Ehlén et al., 2020). In all cases, the BRCA2 fragments became folded upon transient interactions characterized by affinities on the micromolar range.

In this study, we provide a detailed biophysical characterization of the HSF2BP-BRCA2 interaction and the changes in oligomeric state it induces. Its low-nanomolar affinity is orders of magnitude stronger than any other measured interaction involving BRCA2. We also describe the 3D structure of the complex between HSF2BP and BRCA2, which confirms the predicted ARM fold of HSF2BP and reveals the existence of a cryptic repeated motif encoded by exons 12-13 of BRCA2, responsible for binding to ARM oligomers. Finally, using a mouse line engineered to disrupt this high-affinity interaction, we found that contrary to the prediction of the “meiotic localizer” model, the interaction of BRCA2 with HSF2BP is not required for meiotic HR.

## Results

### High affinity interaction between the armadillo domain of HSF2BP and a disordered 52-aa BRCA2 peptide

Our previous analyses using co-immunoprecipitation identified the C-terminal part of HSF2BP (I93-V334) and the BRCA2 fragment G2270-T2337 as interacting regions (Brandsma et al., 2019). Extending this approach (Figure 1A-C, Supplementary Figure S1A), we narrowed down the minimal interaction region to E122-V334 (HSF2BP-H3, hereafter referred as ARM, for “armadillo domain”) and in BRCA2 to N2288-T2337 (fragment F15, **Figure 1B**). Further truncations resulted in loss or reduction in co-precipitation efficiency. We also extended our site-directed mutagenesis mapping: using a homology model of the HSF2BP ARM domain, we predicted structural neighbors of R200, which we previously found to be required for BRCA2 binding, and made substitutions based on human polymorphism data (dbSNP). In contrast to our initial blind screen, most of these substitutions disrupted the HSF2BP-BRCA2 interaction (**Figure 1A**)(Brandsma et al., 2019). To characterize the direct interaction between HSF2BP and BRCA2 *in vitro*, we purified the HSF2BP and ARM proteins, as well as a large region of BRCA2, F0 S2213-Q2342, including F15, that shows high conservation during evolution (**Supplementary Figure S2**). We first performed an NMR analysis of F0, in order to identify the residues binding to ARM. Assignment and further analysis of the NMR Hn, N, Cα, Cβ and Co chemical shifts of ^15^N, ^13^C labeled BRCA2-F0 showed that this peptide is disordered in solution: only region N2291-S2303 forms a transient α-helix present in more than 25% of the molecules (**Supplementary Figure S3**). Addition of unlabeled ARM causes a global decrease of the intensities of the 2D NMR ^1^H-^15^N HSQC peaks of ^15^N labeled F0, with region S2252-Q2342 (further called F_NMR_) showing the largest decrease (intensity ratio lower than 0.4; **Figure 1D**). We concluded that the chemical environment of this region, including F15, is significantly modified in the presence of the ARM domain. Using Isothermal Titration Calorimetry (ITC) experiments revealed that both HSF2BP and ARM bind to F0 with a nanomolar affinity and a stoichiometry of 0.5, i.e. two HSF2BP/ARM bind to one BRCA2 peptide (**Figure 1E; Table 1**). Full-length HSF2BP binds about 25-fold tighter to F0, as compared to ARM. As a control, we verified that HSF2BP mutant R200T does not bind to F0, consistently with our previous report (Brandsma et al., 2019). We further compared the affinity of ARM for F0, F_NMR_ and F15X (the recombinant peptide N2291-Q2342, similar to F15 N2288-T2337 used in cellular assays). The affinities of ARM for F_NMR_ and F15X are not significantly different, being around 10 nM (**Figure 1E; Table 1**). Unexpectedly, they are about 3-fold higher than the affinity measured between ARM and F0 (**Figure 1E; Table 1**). Therefore, we decided to continue by focusing on the complex between ARM and F15X.

**Figure 1.**
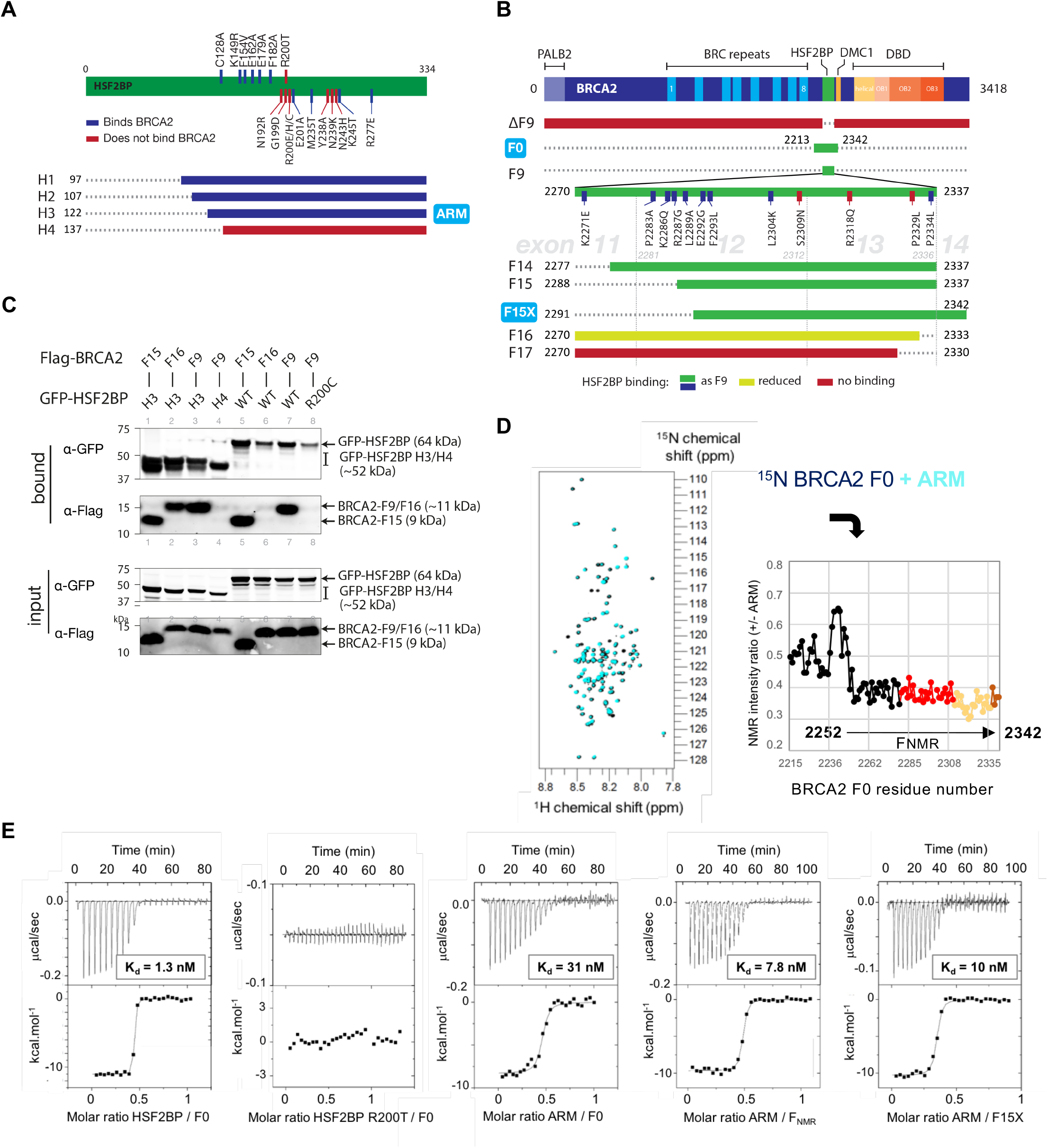
The ARM domain of HSF2BP binds with a nanomolar affinity to a 52 aa BRCA2 peptide. (**A**) Schematic depiction of the truncation and substitution variants of HSF2BP used in this study. Substitutions tested previously are mapped above the bar, whereas those tested in this study are indicated below the bar. Truncation variants are colored based on their ability to bind BRCA2 peptides. (**B**) Schematic depiction of BRCA2 fragments and variants used in the study. Full-length BRCA2 is shown at the top with key domains indicated. Location of the fragment F9 identified previously and its truncations tested here are shown, with colors indicating ability to bind HSF2BP. Fragments produced as recombinant proteins are indicated with blue labels. (**C**) Co-immunoprecipitation of GFP-HSF2BP (full-length wild-type (WT) or R200C variant, and fragments H3 (ARM) and H4) and indicated Flag-tagged BRCA2 variants. Proteins were transiently produced in HEK293T cells. (**D**) NMR characterization of BRCA2 residues involved in binding to the Armadillo domain of HSF2BP (ARM). 2D ^1^H-^15^N HSQC spectra were recorded at 950 MHz and 283 K on the ^15^N-labeled BRCA2 fragment F0 (S2213-Q2342), either free (100 μM; dark blue) or in the presence of the unlabeled ARM domain (1:1 ratio; cyan). Ratios of peak intensities in the two conditions revealed that a set of peaks, corresponding to BRCA2 fragment F_NMR_ (S2252-Q2342), decreased by more than 60% after addition of ARM. The points and curve fragments in red, orange and brown correspond to residues encoded by exon 12, 13 and 14 of BRCA2, respectively. (**E**) ITC curves that reveal how either HSF2BP or its ARM domain (in the instrument cell) interacts with the BRCA2 fragment F0, F_NMR_ and F15X (N2291-Q2342) (in the instrument syringe). The dissociation constants (K_d_) are indicated. All experiments were duplicated, and the dissociation constants, stoichiometry, and thermodynamics parameters of each experiment are recapitulated in Table 1.

**Table 1.** Isothermal Titration Calorimetry data.

| | Kd (± error) (M) | n | $\Delta H$ (±error)<br>(kcal/mol) | $\Delta G$<br>(kcal/mol) | $-T\Delta S$<br>(kcal/mol) |
| --- | --- | --- | --- | --- | --- |
| HSF2BP FL vs F0 - expt 1 | 1.3E-09 (0.3) | 0.43 | -11.0 (0.1) | -11.8 | -0.8 |
| HSF2BP FL vs F0 - expt 2 | 1.3E-09 (0.3) | 0.47 | -10.9 (0.1) | -11.9 | -1.0 |
| HSF2BP FL vs F15XΔ12 - expt 1 | 1.5E-06 (0.7) | 0.89 | -1.7 (0.3) | -7.8 | -6.1 |
| HSF2BP FL vs F15XΔ12 - expt 2 | 2.1E-06 (0.9) | 0.90 | -1.8 (0.2) | -7.6 | -5.8 |
| ARM vs F0 - expt 1 | 31E-09 (9.0) | 0.45 | -8.3 (0.2) | -10 | -1.7 |
| ARM vs F0 - expt 2 | 25E-09 (7.0) | 0.49 | -11.9 (0.2) | -10.1 | 1.8 |
| ARM vs F<sub>NMR</sub> - expt 1 | 7.8E-09 (1.3) | 0.47 | -9.6 (0.1) | -10.8 | -1.2 |
| ARM vs F<sub>NMR</sub> - expt 2 | 13E-09 (4.0) | 0.51 | -9.4 (0.2) | -10.5 | -1.1 |
| ARM vs F15X - expt 1 | 10E-09 (2.1) | 0.33 | -10.2 (0.1) | -10.6 | -0.4 |
| ARM vs F15X - expt 2 | 9.4E-09 (2.3) | 0.40 | -9.2 (0.1) | -10.7 | -1.5 |

### The armadillo domain of HSF2BP tetramerizes upon binding to BRCA2

First, we characterized the molecular mass of the ARM domain either free or bound to F15X. Biophysical analysis by SEC-MALS (Size Exclusion Chromatography - Multiple Angle Light Scattering) and SEC-SAXS (Small Angle X-ray Scattering) revealed that, if free ARM is dimeric, the complex is tetrameric with an estimated molecular weight of 94 kDa (SEC-MALS; **Figure 2A**) or 109 kDa (SEC-SAXS; **Figure 2B**), for a theoretical mass of 4 ARM bound to 2 F15X of 108 kDa. In parallel, intensity curves measured by SAXS on the complex gave a distance distribution reflecting a nearly spherical shape, with a Dmax of 104 Å and a Rg of 34 Å (**Figure 2B**).

**Figure 2.**
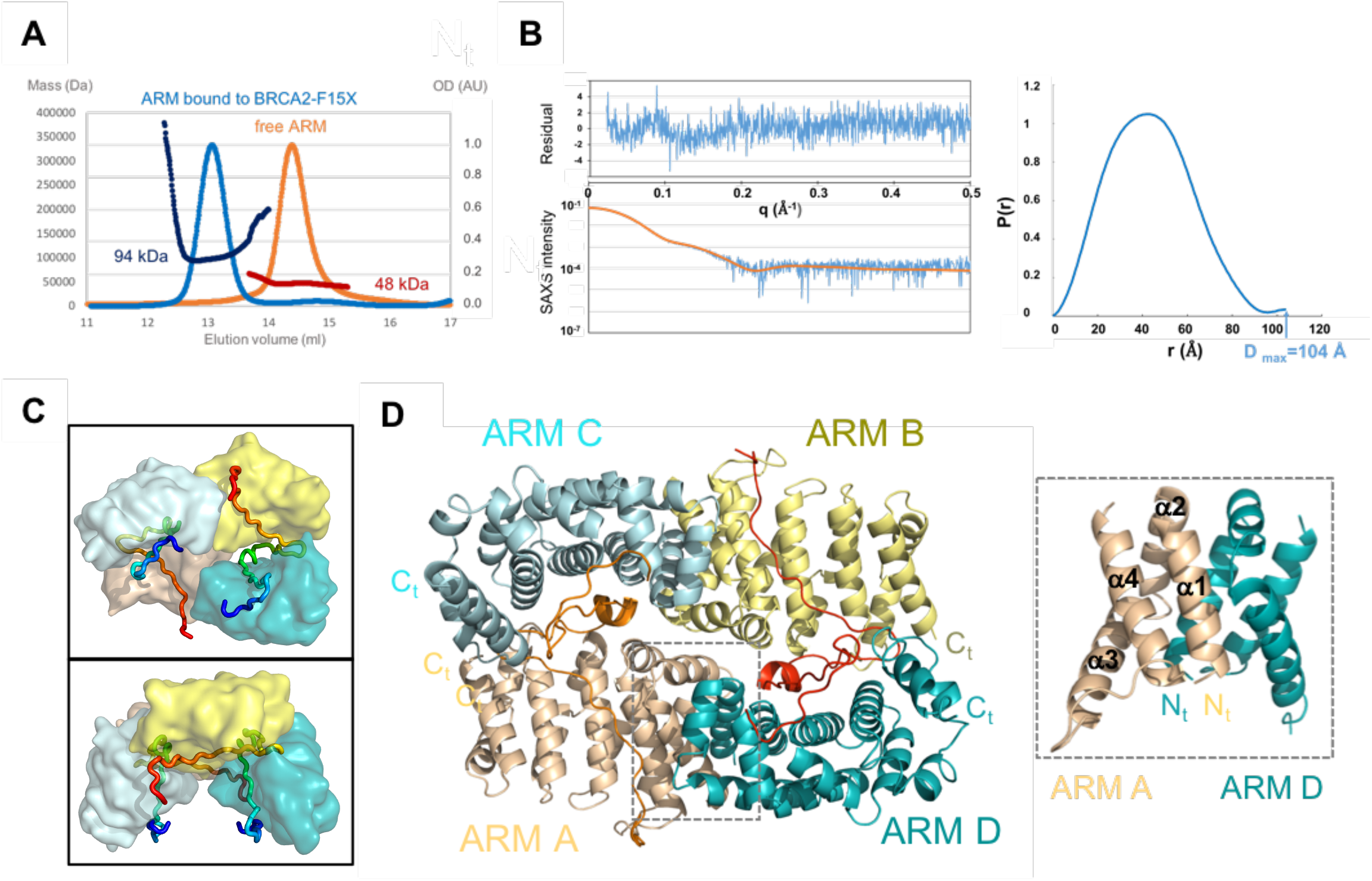
The ARM domain tetramerizes upon BRCA2 binding. **(A)** SEC-MALS profiles from two independent experiments, performed either on free ARM (orange: OD normalized to 1; red: mass) or on ARM bound to F15X (light blue: OD normalized to 1; dark blue: mass) (column: Superdex 200 10/300 GL). **(B)** SEC-SAXS curve and resulting distance distribution obtained on ARM bound to F15X (blue). The experimental SAXS curve is compared to the theoretical SAXS curve calculated with CRYSOL from the X-ray structure of the complex (orange). Residual errors are plotted as a function of the scattering angle (resulting chi^2^ value: 1.8 Å^2^). **(C, D)** Different views of the crystal structure of the complex, illustrating how the ARM dimers, formed by chains A (wheat) and D (teal) and chains B (yellow) and C (pale blue), are held together through their interactions with the BRCA2 peptides. In (**C**), the ARM domains are represented as surfaces, whereas the BRCA2 peptides are displayed as tubes colored from their N-terminus (blue) to their C-terminus (red). In (**D**), all chains are represented as cartoons, the BRCA2 peptides being colored in orange (chain E) and red (chain F). A zoom view of the dimerization interface between chains A and D is displayed in a dotted box.

Crystals of the complex were obtained within a few days by hanging-drop vapor diffusion and diffracted up to 2.6 Å on PROXIMA-1 and PROXIMA-2 beamlines at the SOLEIL synchrotron. The structure of the complex was solved using a combination of Molecular Replacement and SAD approaches (see details in Material and Methods; **Table 2** and **Supplementary Figure S4**). The overall conformation of the structure is consistent with the SAXS data obtained in solution, as reflected by the low chi^2^ value of 1.8 Å^2^ obtained when fitting the SAXS curve deduced from the experimental structure to the experimental SAXS curve (**Figure 2B**).

**Table 2.**
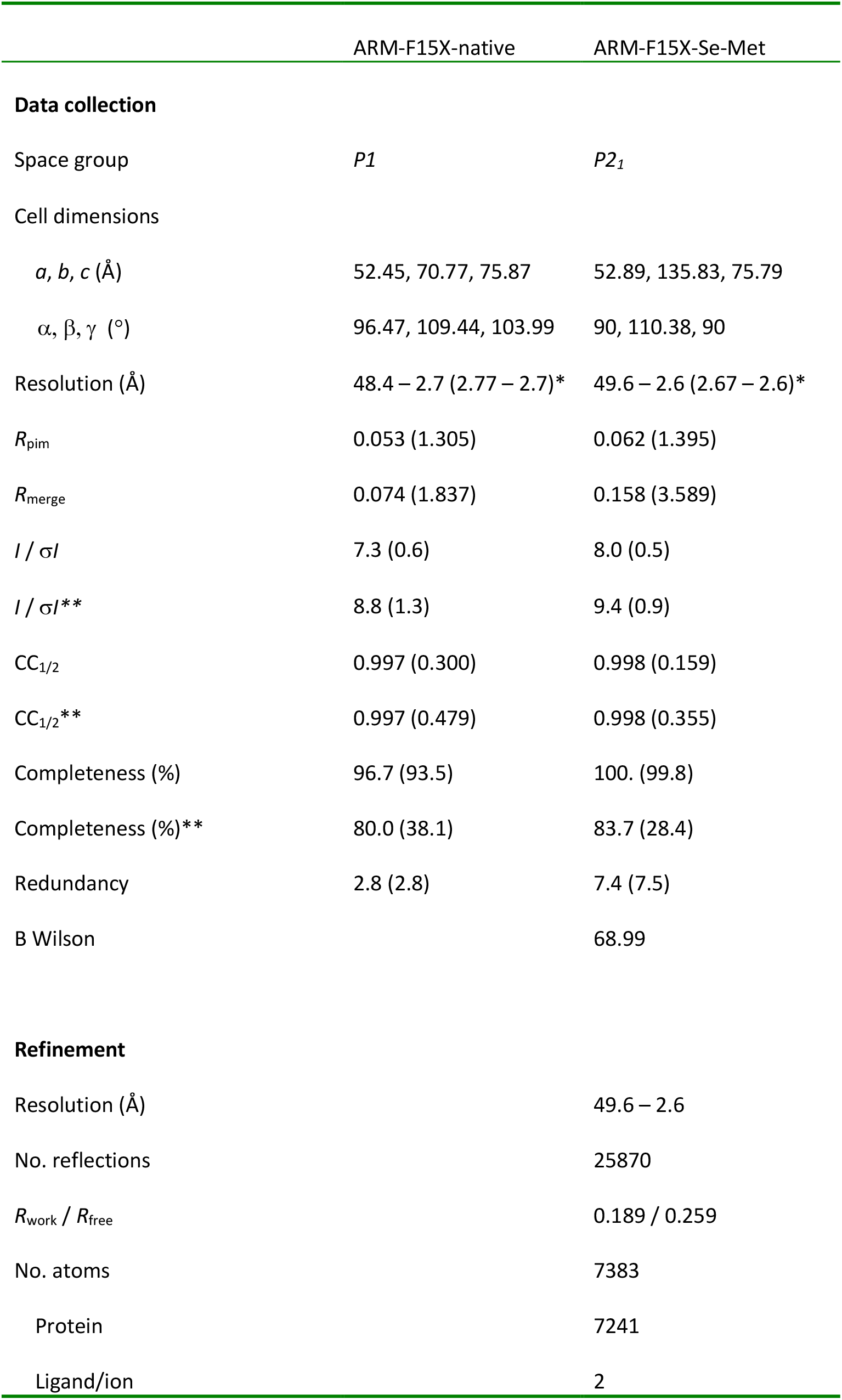
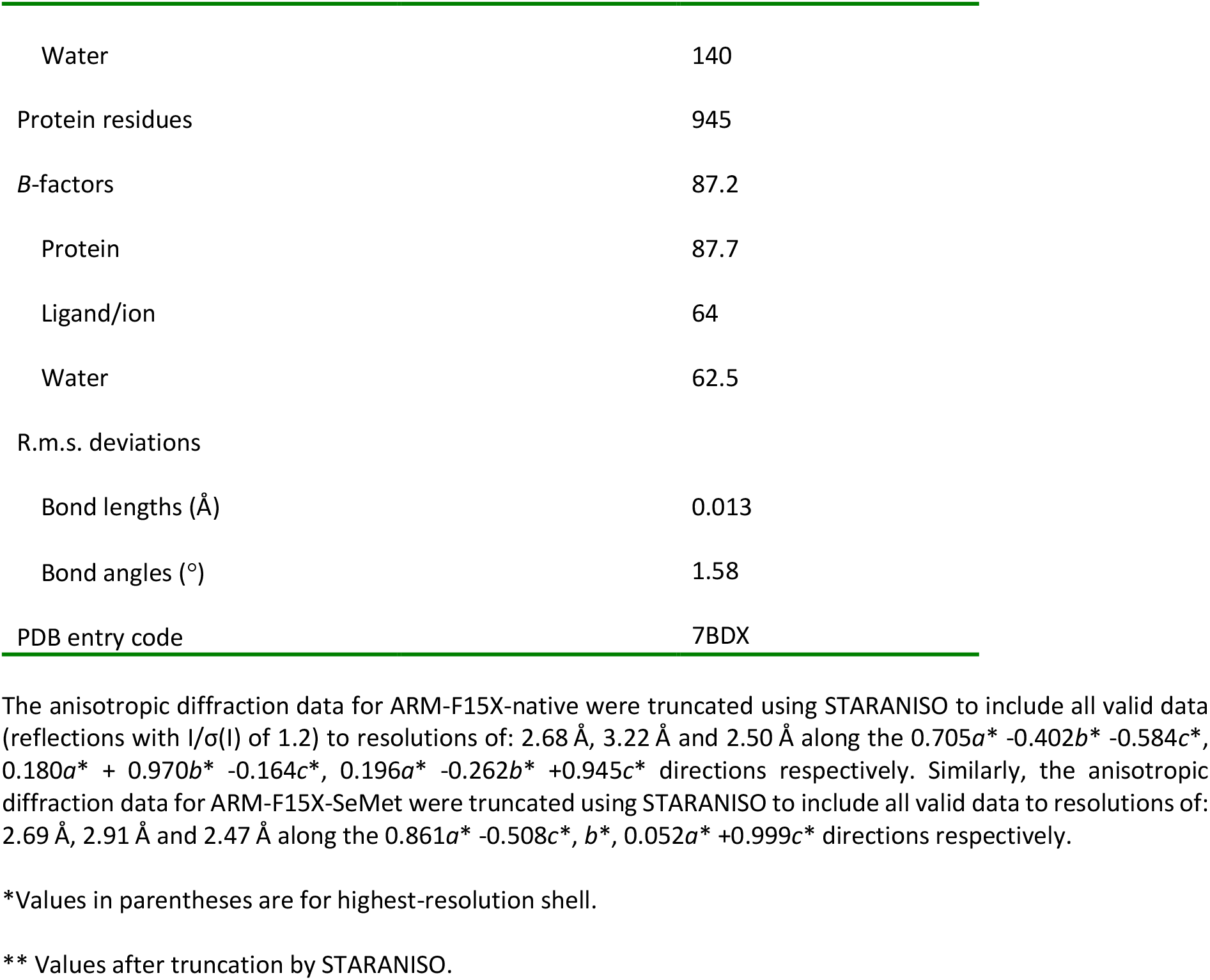
Data collection and refinement statistics.

The crystal structure includes four ARM domains and two BRCA2 F15X peptides (**Figure 2C,D**). The ARM domains A and D, as well as B and C, form dimers through a symmetric interface of about 950 Å^2^. This interface involves their N-terminal fragments from E122 to I156 (**Figure 2D**; **Supplementary Figure S5A**); it is mediated by hydrophobic residues from helices a1 (M123, A126, A127, L130, L131, V134), helices α2 (V140, I144) and the N-terminus of helices α3 (L151, F153, I156); about a quarter of this interface is due to the interaction between the highly conserved L131 and F153 from one monomer and the highly conserved L130 from the other monomer. The ARM domains A and C, as well as B and D, have a very small direct interface of less than 100 Å^2^. They are juxtaposed, one chain being rotated around its main axis by about 90° relatively to the other, and interact mainly through BRCA2 (**Figure 2C**). The BRCA2 peptide in orange (chain E) runs along the V shaped groove formed by chains A and C. Similarly, the other BRCA2 peptide (in red; chain F) runs along the groove formed by chains B and D. At the center of the tetramer, a symmetric interface of about 250 Å^2^ is observed between the ARM domains A and B, which involves helices α2 (K142, A143, K149 and A150) and helices α5 (N205, S206) (**Supplementary Figure S5B**). This interface is poorly conserved. In summary, two ARM dimers interact through two BRCA2 peptides to form a tetramer; within the tetramer, two types of conserved interfaces are observed, either between monomers from the same dimer (chains A and D, as well as B and C), or between the ARM domains and the BRCA2 peptides.

### Two ARM dimers are held together by two BRCA2 peptides through repeated motifs

The 3D structures of the complexes between, on the one hand, the ARM domains A and C and the peptide E, and on the other hand the ARM domains B and D and the peptide F, are remarkably similar (**Figure 3A**). In these structures, two ARM monomers form a BRCA2-binding surface of 2740 Å^2^, which is in the upper range of interaction surfaces, even for a complex between a folded domain and an intrinsically disordered peptide (Mészáros et al., 2007). The BRCA2 peptide engages 48 aa in this interaction. The ARM domains C and D interact with the N-terminal sequence of the BRCA2 peptides, from N2291 to E2328, whereas the ARM domains A and B interact with their C-terminal sequence, from D2312 to T2338. The 3D structures of the ARM domains interacting with the same region of the peptide are highly similar, whereas the 3D structures of two ARM domains interacting with different regions of the BRCA2 peptide show some local structural variations, as measured by the root-mean-square deviations between their Cα atoms (see table in **Figure 3A** and **Supplementary Figure S6**). Another remarkable feature of this complex is that the same surface of the ARM domains recognizes the N-terminal and the C-terminal regions of the BRCA2 peptide (**Figure 3B; Supplementary Figure S7**). Indeed, a surface of about 1540 Å^2^ formed by helix α1, helix α4 and the N-terminal region of α5, helix α7 and the N-terminal region of α8, helix α10 and loop α10α11, and finally loop α12α13 on one ARM domain interacts with the N-terminal sequence of the BRCA2 peptide. A smaller surface of 1200 Å^2^ formed by helix α1, helix α4 and the N-terminal region of α5, helix α7 and the N-terminal region of α8, only the C-terminal region of α10 and loop α10α11 on the other ARM domain interacts with the C-terminal sequence of the peptide. The surface common to the two binding interfaces is conserved through evolution and positively charged (**Figure 3C**). The surface specific to the interface with the N-terminal region of the peptide, including the N-terminus of helices α7 and α10 and loop α12α13, is less conserved. It contacts the central region of the BRCA2 peptide, from I2315 to E2328 of BRCA2, in particular through a set of hydrogen bonds and salt bridges between ARM D268, E270 and BRCA2 R3219, and between ARM R315 and BRCA2 S2326, E2328.

**Figure 3.**
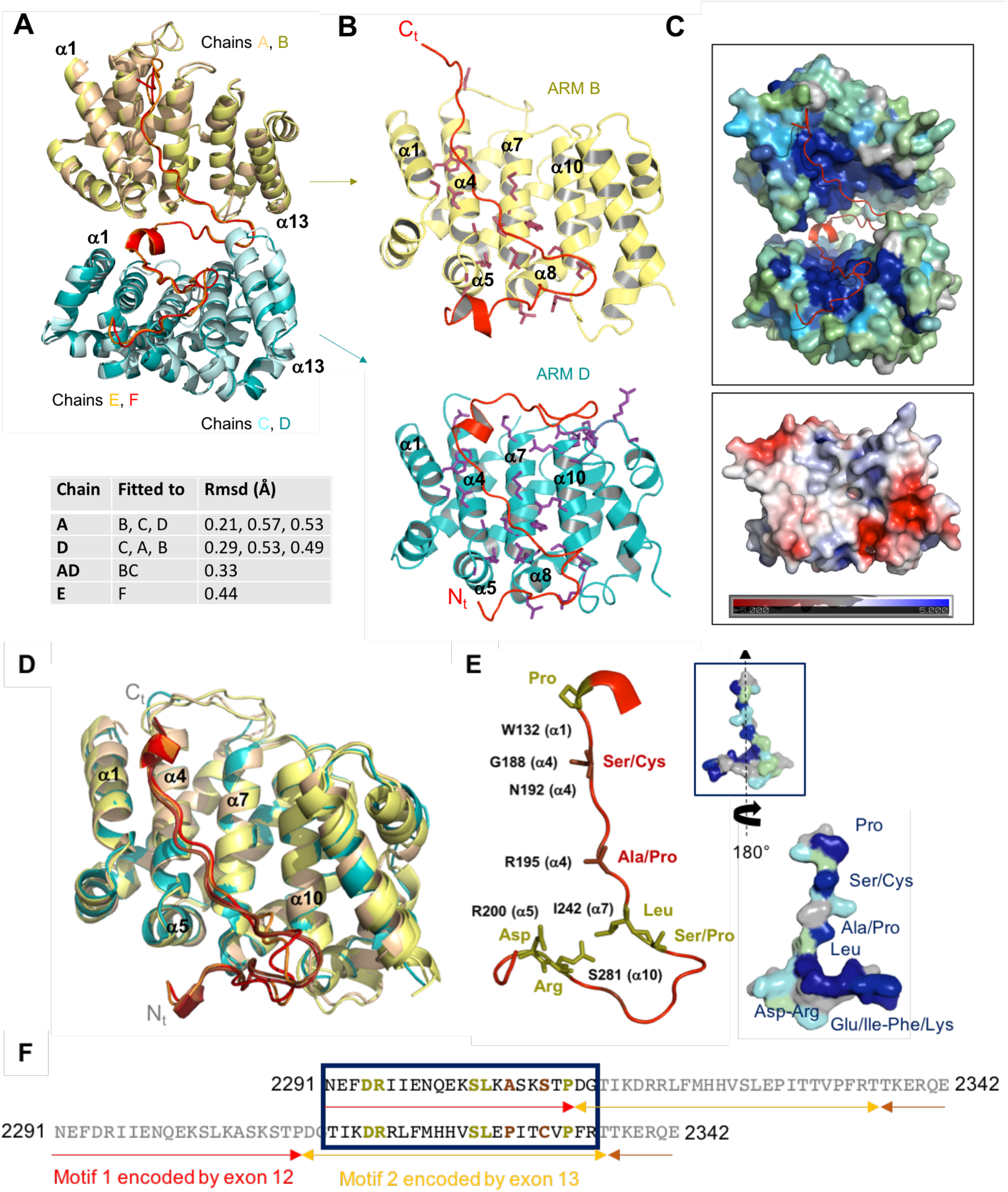
The same conserved ARM surface interacts with both the N-terminal and C-terminal regions of the BRCA2 peptide, through a new 23 aa motif. **(A)** Superimposition of the complexes formed by two ARM domains and a BRCA2 peptide. The Ca root-mean-square-deviation (Rmsd) values calculated between the different chains are recapitulated in the lower part of the panel. Additional views of the superimposed individual chains are displayed in Supplementary Figure S7. **(B)** Zoom on the interfaces between chain B and the C-terminal region of F (upper view) and chain D and the N-terminal region of F (lower view). ARM residues that are either involved in hydrogen bonds or salt bridges with BRCA2, or buried (by more than 30 Å^2^) at the interface with BRCA2, are represented by colored sticks. They are labeled in Supplementary Figure S7B,C. **(C)** Surfaces of ARM domains, colored as a function of sequence conservation (upper panel) or electrostatic potential (lower panel). In the upper panel, surfaces of chains B and D are colored as a function of conservation scores calculated by Consurf (Ashkenazy et al., NAR 2016). High, medium, weak and no conservation are indicated in dark blue, cyan, green and grey, respectively. Chain F is represented as a red ribbon. In the lower panel, only chain D is displayed, and its surface is colored from red (negatively charged) to blue (positively charged). **(D)** Superimposition of the 4 ARM domains and their BRCA2 interacting peptides. The 4 ARM structures were aligned, and their BRCA2 interacting fragments are displayed. **(E)** Representation of the 3D structure of the BRCA2 motif binding to ARM domains. Residues strictly conserved between motif 1 and motif 2 are displayed as olive sticks. They correspond to D2294/2317, R2295/2318, S2303/2326, L2304/2327, P2311/2334. Residues that are similar between the two motifs and conserved in BRCA2 are displayed as brown sticks. They correspond to A2306/P2329 and S2309/C2332 (mutated in T2332 in our construct to avoid oxidation of this residue). A set of conserved residues from ARM interacting with BRCA2, as defined in (B), are indicated in black next to the BRCA2 residues when they directly interact with these residues. A boxed surface view of the peptide, in the same orientation as the cartoon view, is colored by conservation as in (C). Turning this surface view by 180°reveals the conservation of the surface binding to ARM, including the conserved residues of motifs 1 and 2. **(F)** Sequence alignment of motif 1, interacting with chains C and D, and motif 2, interacting with A and B. Conserved residues are colored as in (E). Motif 1 is encoded by exon 12, whereas motif 2 is encoded by exon 13.

Further analysis of the interface revealed that, even more strikingly, the N-terminal and C-terminal sequences of the BRCA2 peptide interacting with the ARM domains have similar structures (**Figure 3D**). BRCA2 motif 1, from N2291 to G2313, and BRCA2 motif 2, from T2314 to R2336, can be nicely superimposed, and interact with the same ARM surfaces. Sequence alignment between the two motifs identified 5 identical residues: D2294/2317, R2295/2318, S2303/2326 (also a proline in some organisms), L2304/2327 and P2311/2334, and 2 similar residues: A2306/P2329, S2309/C2332 (T2332 in our construct), which are all conserved through evolution (**Figure 3E; Supplementary Figure S2**). These residues interact with conserved residues from the ARM domains, as indicated in **Figure 3E**. For example, D2294/2317 contacts R200 (helix a5) through a set of hydrogen bonds/salt bridges; R2295/2318 is hydrogen-bonded to S281 (helix a10); L2304/2327 contacts I242 (helix a7); S2309/C2332 and P2311/2334 interact W132 (helix α1); S2309/C2332 also contacts G188 and is hydrogen-bonded to N192 (helix α4). These conserved interactions between motif 1 and motif 2 of BRCA2 and two ARM monomers belonging to two different dimers triggers tetramerization of the ARM domain. In summary, analysis of the crystal structure of the BRCA2-HSF2BP complex identified a new repeated motif in BRCA2 that is able to bind to the ARM domain of HSF2BP, thus causing tetramerization of the dimeric ARM domain.

### Critical residues identified in the structure are essential for HSF2BP binding to BRCA2 in cells

We compared the interface observed in our crystal structure with the point mutants we analyzed by co-immunoprecipitation (**Figure 1A-C**). Substitutions N192R, R200E/H/C, M235T, Y238A, N239K affected residues involved in binding to both motifs 1 and 2: most of them totally abolished binding, only M235T still weakly bound to BRCA2 (**Figure 4A; Supplementary Figure S1**). Substitutions G199D, E201A, N243H and R277E affected additional residues that are conserved and partially buried due to BRCA2 binding: both G119D and N243H abolished binding, and R277E strongly decreased binding (**Figure 4A; Supplementary Figure S1**). In contrast, E201A and K245T affected two less-conserved residues, and these mutations had a weak (or not observable) impact on BRCA2 binding (**Figure 4A**). Altogether, co-immunoprecipitation assays validated the essential role played by the conserved ARM surface in binding to BRCA2 in human cells. Furthermore, these assays highlighted the critical role played by asparagine residues from the ARM surface. Despite the lack of secondary structure elements in bound BRCA2, the presence of asparagine residues that interact through their side chains with the backbone amide proton and oxygen of BRCA2 residues mimics the formation of a β-sheet hydrogen bond network between the ARM domains and BRCA2 (**Figure 4B**). These interactions are independent of the BRCA2 sequence, as they involve only the backbone atoms of BRCA2. On the BRCA2 side, consistently with our ITC results, the only mutations clearly decreasing or abolishing the interaction are in the region L2304-P2329 (**Figure 1B**). Most mutations of residues defining motif 1 (L2304K, S2309N), and motif 2 (R2318Q, P2329L) either strongly decreased or abolished binding to HSF2BP (**Figure 4C**). Only P2334L that also affected a residue defining motif 2, did not impair binding to HSF2BP. More generally, mutations of the highly conserved BRCA2 residues E2292, F2293 and P2334 did not result in any loss of binding, despite F2293 and P2334 being buried at the interface with HSF2BP. However, these residues were mutated into leucine: their hydrophobic character was conserved, which might explain the lack of associated binding defect. In summary, the impact of human polymorphisms at the HSF2BP-BRCA2 interface was characterized, and a set of mutations of highly conserved residues was identified that severely decreased the interaction between these two proteins in cells.

**Figure 4.**
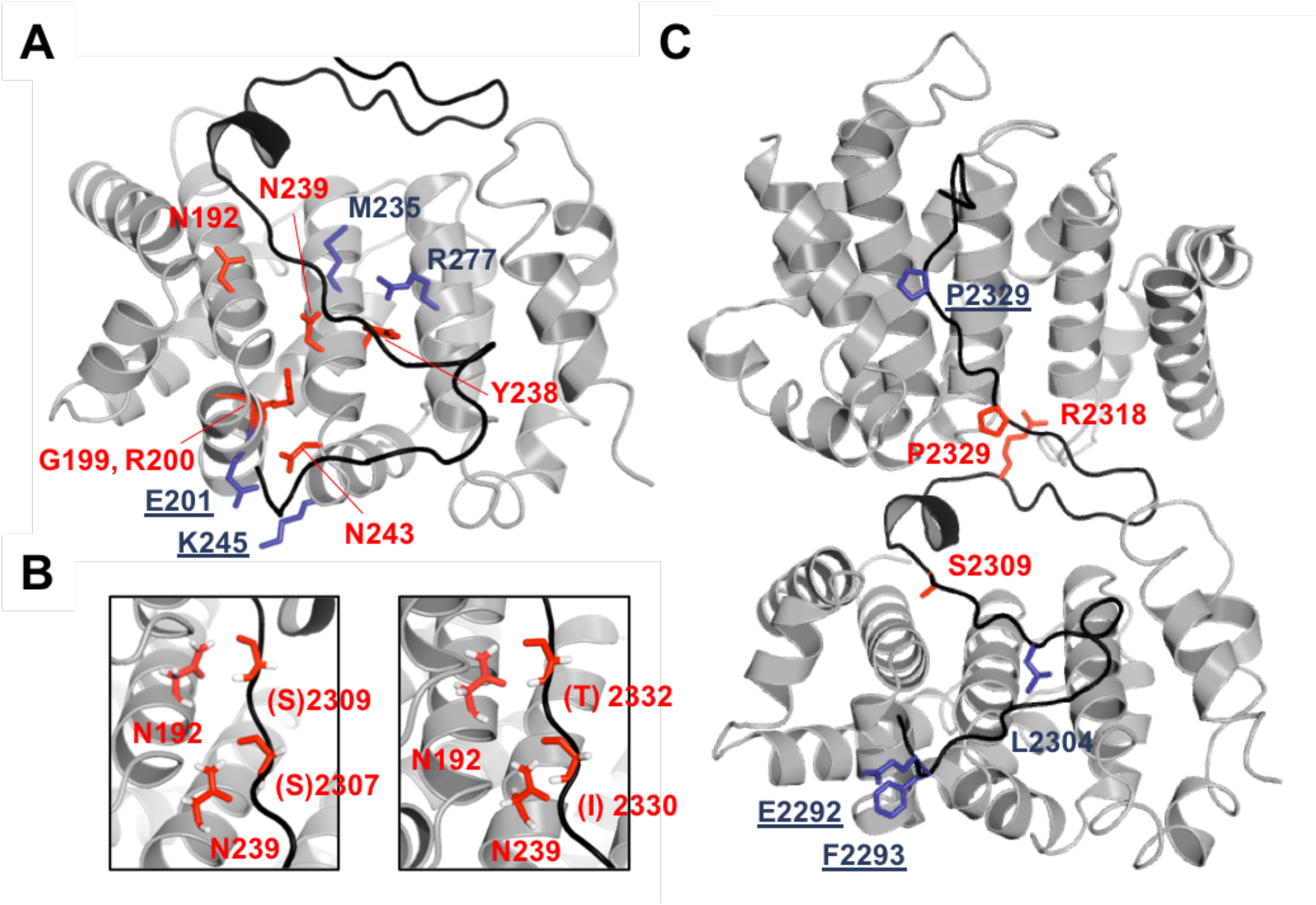
Screening of the interface by mutagenesis identifies mutations abolishing binding. **(A)** 3D view of the ARM chain D in complex with motif 1 of the BRCA2 chain F. The ARM and BRCA2 protein fragments are displayed as grey and black ribbons, respectively. The side chains substituted for coimmunoprecipitation studies are displayed in sticks, and colored as in Figure 1A (red: no binding; blue: residual binding with the residue name underlined when wild-type binding was observed). **(B)** Zoom views on residues forming a hydrogen bond network between ARM and BRCA2. Asparagine side chains from ARM interact with backbone atoms from BRCA2. The side chains of BRCA2 are not displayed; consistently, the BRCA2 residue names are in brackets. Hydrogens were added to the crystal structure for clarity. **(C)** 3D view of the ARM chains B and D in complex with the BRCA2 chain F. The ARM and BRCA2 protein fragments, as well as the mutated side chains, are displayed as in (**A**), except that full labels are shown for BRCA2 residues.

### Deletion of Brca2 exon 12 disrupts functional interaction between BRCA2 and HSF2BP in cells

Previously we showed that excising exon 12 from *BRCA2* in human cells, mimicking a naturally occurring *BRCA2* splice form, renders them completely resistant to the inhibitory effect HSF2BP has on HR in the context of DNA inter-strand crosslink repair (Sato et al., 2020). This suggested complete disruption of the functional interaction between HSF2BP and BRCA2Δ12. Analysis of our crystal structure revealed that exon 12 encodes motif 1, whereas exon 13 encodes motif 2 (**Figure 5A**). Thus, deleting exon 12 should lead to the loss of more than half of the binding interface, and also to the absence of resulting tetramerization of the ARM domain. To test this hypothesis, we measured the affinity of HSF2BP for the F15X peptide deleted from the sequence encoded by exon 12, named F15XΔ12 (D2312-E2342). This affinity is on the micromolar range, that is 1000-fold weaker than the affinity of HSF2BP for F0 (**Figure 5B**). Moreover, the stoichiometry of the interaction is now 1, demonstrating that each HSF2BP molecule binds to its own BRCA2 peptide. Further analysis of the ARM-F15XΔ12 complex using SEC consistently showed that the ARM domain does not oligomerize upon binding to F15XΔ12 (**Figure 5C**). To test the effect of exon 12 loss under physiological conditions, we created the *Brca2* exon 12 deletion in mouse embryonic stem (ES) cells, where HSF2BP is expressed natively (Brandsma et al., 2019). To further validate the specificity of our system, we also engineered another *Brca2* in-frame exon excision, deleting exons 12-14 which encode all of BRCA2 residues involved in the interaction with HSF2BP (**Figure 5D**). In addition to the different *Brca2* exon excisions we homozygously knocked-in *GFP* expression sequence at the 3’ end of *Hsf2bp* or *Brca2* coding sequence (Reuter 2014, Brandsma 2019). This allowed us to study HSF2BP-BRCA2 interactions under native expression levels, and at the same time take advantage of the highly efficient GFP nanobody precipitation, thus reducing the chance of missing any possible residual interaction between the proteins, while minimizing non-specific background associated with indirect immunoprecipitation. To avoid non-linear amplification in immunoblotting, we used fluorescently labeled secondary antibodies instead of enzymatic detection. Pull-down from cells producing full-length, Δ12 or Δ12-14 BRCA2-GFP from engineered homozygous alleles revealed near-complete (by 95±3%, n=4) and complete abrogation of HSF2BP co-precipitation in Δ12 and Δ12-14, respectively (**Figure 5D**). Pull-down of HSF2BP-GFP from *Hsf2bp^GFP/GFP^Brca2^wt/wt OR Δ12/Δ12 OR Δ12-14/Δ12-14^* ES cells revealed somewhat higher disruption (97-99%, n=2) of co-precipitation in BRCA2-Δ12, and only background signal for BRCA2-Δ12-14 (**Figure 5E**). We also tested co-precipitation between human proteins in HeLa cells overproducing GFP-HSF2BP and producing full-length, Δ12 or Δ12-14 BRCA2 from engineered native alleles. Human BRCA2 and HSF2BP behaved similar to the mouse proteins (**Figure 5F**) and consistent with the functional experiments we described before (Sato 2020). Thus, in agreement with our biophysical data, loss of *Brca2* exon 12 strongly decreased interaction with HSF2BP. To evaluate its effect on HSF2BP-BRCA2 in functional contexts in cells, we analyzed HSF2BP-GFP diffusion in living cells and its recruitment to ionizing radiation induced nuclear foci, using the same engineered *Hsf2bp* and *Brca2* allele combinations in mES cells. Characteristic (BRCA2-like) constrained diffusion of HSF2BP-GFP we described before (Brandsma 2019) was dramatically affected by exons 12-14 deletions; in particular the slow-diffusing and immobile species were gone (**Supplementary Movies 1-3**). We further noted that in *Brca2* Δ12 and Δ12-14 cells, HSF2BP-GFP fluorescence intensity in the nucleus was reduced, and more fluorescence was observed in the cytoplasm, which made it altogether impossible to apply the quantitative single particle tracking analysis we used before. We observed a similar reduction in nuclear fluorescence and co-localization of HSF2BP-GFP with RAD51 in ionizing radiation induced nuclear foci in immunofluorescence experiments (**Figure 5G,H**). Taken together, this and the functional experiments in human *BRCA2^Δ12/Δ12^* cells, indicate that exclusion of the BRCA2 domain encoded by exon 12 leads to a severe defect in the interaction with HSF2BP in cells.

**Figure 5.**
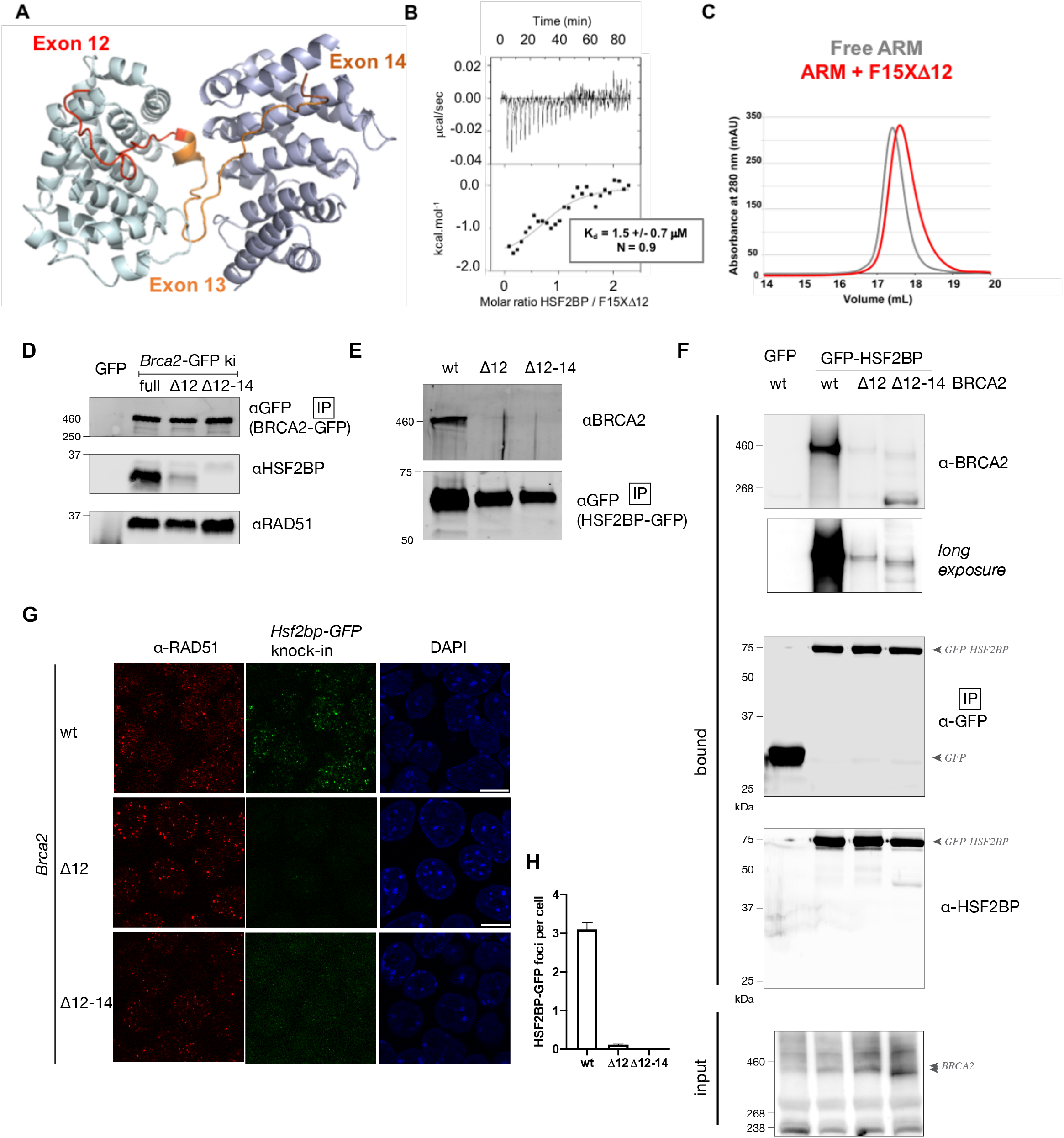
Deletion of *Brca2* exon 12 disrupts HSF2BP-BRCA2 interaction in cells. (**A**) Structure of the HSF2BP dimer interacting with the BRCA2 peptide colored by encoding exon number. (**B**) ITC experiment with HSF2BP and a truncated variant of F15X peptide missing residues encoded by exon 12 (F15XΔ12). (**C**) ARM and the complex between ARM and F15XΔ12 analyzed by analytical gel filtration (column: Superose 6 Increase 10/300 GL) and SDS-PAGE (data not shown). (**D**) Immunoblot analysis of proteins co-precipitated with anti-GFP nanobody beads from mES cells containing homozygous *Brca2-GFP* allele without (full) or with deletions of exon 12 or exons 12-14. (**E**) Immunoblot analysis of proteins co-precipitated with anti-GFP nanobody beads from double knock-in mES cells containing homozygous *Hsf2bp-GFP* and *Brca2* alleles without or with deletion of exons 12 or 12-14. (**F**) Immunoblot analysis of proteins co-precipitated with anti-GFP nanobody beads from HeLa cells stably overproducing GFP-HSF2BP or GFP control, and in which *BRCA2* allele was modified by excision of exons 12 or exons 12-14, or unmodified wild type (wt) *BRCA2*. (**G**) *Hsf2bp^GFP/GFP^Brca2^wt/wt^, Hsf2bp^GFP/GFP^Brca2^Δ12/Δ12^* and *Hsf2bp^GFP/GFP^Brca2^Δ12-14/Δ12-14^* mES cells were irradiated with 8 Gy, fixed after 2h recovery, immunostained with anti-RAD51 antibody, mounted with DAPI, and imaged using laser confocal microscope. HSF2BP-GFP was detected by direct fluorescence. Maximum projection of 3 confocal slices (0.5 μm apart) is shown. (**H**) Quantification of HSF2BP-GF foci from the experiment shown in panel (G).

### High-affinity HSF2BP-BRCA2 interaction is not required for meiotic HR

Our biophysical data and functional experiments validated Brca2-Δ12 as a model to determine the functional importance of the high-affinity interaction between BRCA2 and HSF2BP and of the resulting HSF2BP oligomerization. To study this in the context of meiotic HR, we created a *Brca2^Δ12/Δ12^* mouse model (**Figure 6A,B**). Consistent with the robust proliferation of the engineered BRCA2 Δ12 HeLa and mES cells, *Brca2^Δ12/Δ12^* mice were viable, born at Mendelian ratios, and did not show any overt phenotypes. Moreover, contrary to our expectation, the Δ12 mutation did not phenocopy the *Hsf2bp* knock-out, as not only females, but also males were fertile, with no abnormalities in testes size and sperm counts (**Figure 6C-D**). Morphology of testis tubule sections was also indistinguishable from wild type (**Supplemental Figure S8A**). Molecular analysis of the meiotic prophase progression did not reveal any defects: neither the number of DMC1 and RAD51 foci, nor the timing of their formation in the meiotic prophase I were affected by the mutation we introduced. In addition, the frequency of MLH1 foci indicating meiotic crossover sites after successful HR was normal (**Figure 6E-H, Supplemental Figure S8B**). These results were not consistent with the proposed role of HSF2BP as a BRCA2 localizer in meiosis. We hypothesized that given the high affinity interaction revealed by our ITC experiments, the dependence may be opposite: BRCA2 may bring HSF2BP to the double strand breaks (DSBs). However, the recruitment of HSF2BP and BRME1 to meiotic DSBs was also not affected by the deletion of exon 12 (**Figure 6I-K**).

**Figure 6.**
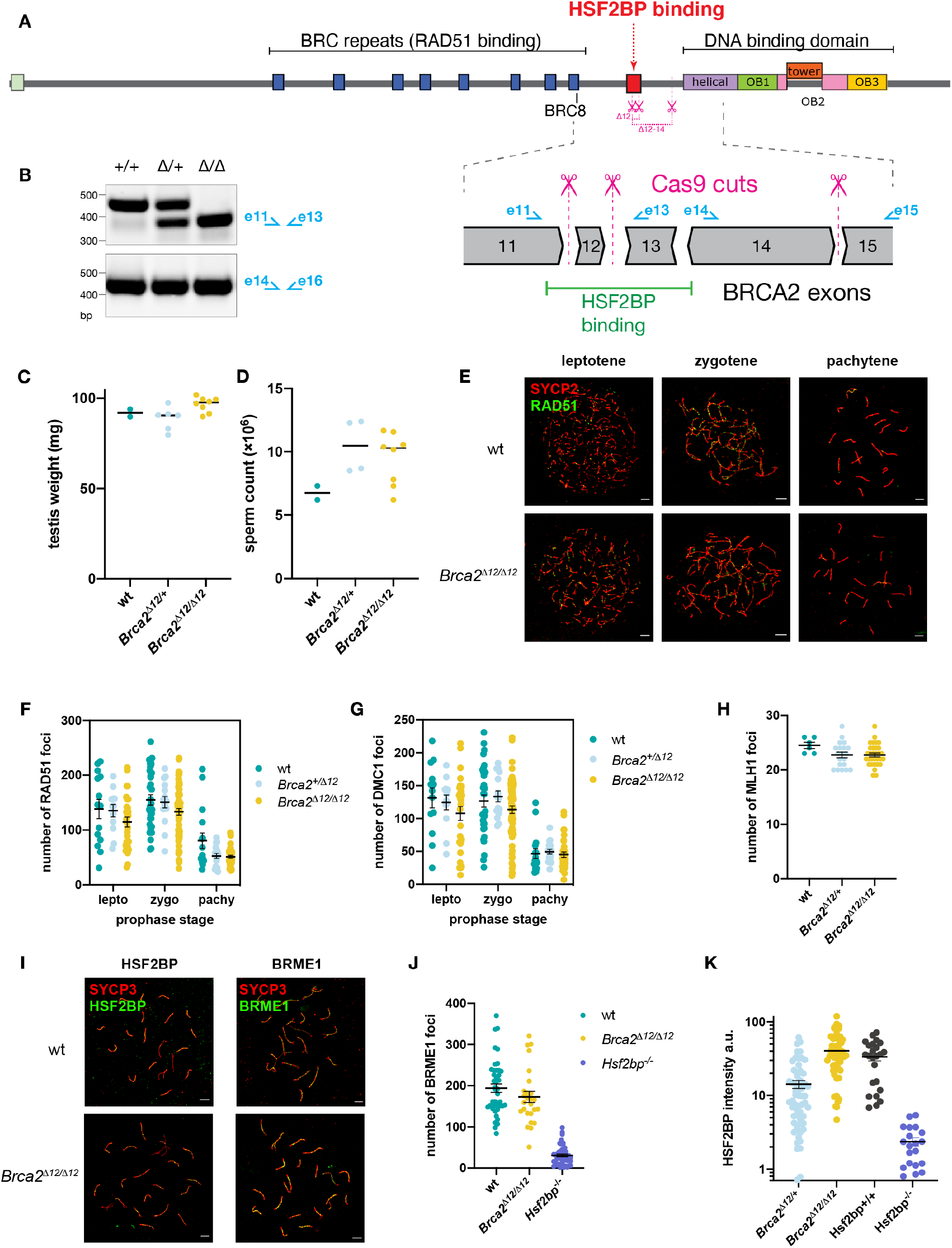
Meiotic phenotype of *Brca2* exon 12 deletion mouse model. **(A)** Schematic depiction of the domain composition of the BRCA2 protein and the exons 11-15 encoding HSF2BP-binding and DMC1-binding domains. Introns are not drawn to scale; different exon phases are indicated by the shape of the boundary. Location of Cas9 cut sites for exon 12 and exons 12-14 excision is shown. (**B**) RT-PCR on cDNA from mouse testis with indicated genotypes confirming the loss of exon 12; primer locations are shown on the exon scheme in panel (A). (**C**) testis weight, (**D**) and sperm count in *Brca2^Δ12/Δ12^* and control mice. (**E-K**) Immunofluorescent analysis of meiotic protein localization on spermatocyte spreads from *Brca2^Δ12/Δ12^* and control mice. Representative images (E,I) and quantification of RAD51 (F), DMC1 (G), MLH1 (H), BRME1 (J) and HSF2BP (K) are shown.

## Discussion

In this paper, we analyzed the structural properties and functional consequences of the BRCA2-HSF2BP interaction and tested the emerging model of its involvement in meiosis. The essential roles of BRCA2 and HSF2BP in meiotic HR have been clearly demonstrated previously. BRCA2 interacts with both RAD51 and DMC1 recombinases, is required for their accumulation at meiotic DSB in mice and stimulates their activity *in vitro*. But how BRCA2 balances its activity with respect to RAD51 and DMC1, and integrates with the meiotic-specific HR machinery in a timely manner during meiosis remains unclear. Direct data on its behavior in meiocytes is scarce, and mechanistic models are mostly based on extrapolation. The proposed role of the recently identified HSF2BP, required for RAD51 and DMC1 accumulation at meiotic DSBs during spermatogenesis in mice (Brandsma et al., 2019), is to bring BRCA2 to meiotic DSBs (Zhang et al., 2019). We tested this hypothesis by eliminating the HSF2BP-binding region of BRCA2 in mice.

We first characterized the BRCA2-HSF2BP interaction *in vitro*. We had previously identified that the region of BRCA2 binding to HSF2BP is located between its BRC repeats and its C-terminal DNA binding domain (**Figure 1B**; (Brandsma et al., 2019)). Our structural analysis unexpectedly revealed that this BRCA2 region contains a duplicated motif, each motif binding to the same residues of an Armadillo domain of HSF2BP (**Figure 3E**). By itself, the Armadillo domain dimerizes through a conserved surface formed by helices α1 to α3 (**Figure 2D**), and presents on each monomer a large conserved groove, which indicates a binding site for functionally important partners (**Figure 3C**). Many Armadillo domains interact through their concave surface with largely disordered partners (Gul et al., 2017). The Armadillo domain of HSF2BP contains 4 Armadillo repeats (**Supplementary Figure S7A**). Altogether, they form a positively charged groove delimitated by helices α1, α4, α7, α10 and α13. This groove is able to recognize a 23 aa motif located in a conserved and disordered region of BRCA2. Because this motif is duplicated in BRCA2, and each motif binds to a different ARM dimer, the interaction triggers further oligomerization of the Armadillo domain into a tetramer. The affinity of BRCA2 for HSF2BP yields 1 nM, which is significantly higher than the affinities yet measured between BRCA2 disordered regions and its partners PALB2, RAD51 and PLK1, all comprised between 100 and 1000 nM (Ehlén et al., 2020; Oliver et al., 2009; Pellegrini et al., 2002). However, after deleting motif 1 encoded by exon 12, motif 2 alone binds to HSF2BP with a micromolar affinity and is unable by itself to trigger oligomerization of the ARM domain into a tetramer (**Figure 5B**).

Consistently with our *in vitro* study, we observed in mouse ES cells that deletion of exon 12, coding for motif 1, causes a severe decrease in the BRCA2-HSF2BP interaction, and deletion of exons 12 to 14, coding for motifs 1 and 2, completely abolishes this interaction (**Figure 5D,E**). We previously showed that HSF2BP mutation R200T abolishes localization of the GFP-tagged HSF2BP protein to mitomycin C-induced repair foci (Brandsma et al., 2019). Consistently, we now demonstrated that the BRCA2 region encoded by exon 12 is responsible for HSF2BP localization at irradiation-induced DSBs (**Figure 5G**). Finally, we previously reported that, in human cells, excising exon 12 from *BRCA2*, mimicking a naturally occurring *BRCA2* splice form, rendered them completely resistant to the inhibitory effect HSF2BP has on HR in the context of DNA inter-strand crosslink repair (Sato et al., 2020).

Based on these structural, biochemical and functional experiments, we developed a *Brca2^Δ12^* mouse line to test the emerging model, which posits that HSF2BP-BRME1 complex acts as a meiotic localizer of BRCA2 (Zhang et al., 2020, 2019), and thus predicts that disengaging BRCA2 from HSF2BP will phenocopy *Hsf2bp* deficiency. However, we could not detect any differences in meiosis in *Brca2^Δ12/Δ12^* mice compared to Brca2^+/+^ (**Figure 6**). Not only females, but also males were fertile. Also, the MLH1 foci indicating meiotic crossover sites after successful HR as well as the DMC1 and RAD51 foci were normal. This showed that the high affinity interaction between BRCA2 and HSF2BP, together with the oligomerisation of HSF2BP triggered by this interaction, are not essential for HR in meiosis.

The BRCA2 localizer function of HSF2BP was suggested primarily based on the observation of recombinant GFP-tagged BRCA2 fragments produced by electroporation of expression constructs into wild-type and HSF2BP-deficient testis (Zhang et al., 2019). In these experiments, a BRCA2 fragment including the HSF2BP binding region and the C-terminal ssDNA binding domain co-localized with RPA2 at DSB sites in an HSF2BP-dependent manner (Zhang et al., 2019). The presence of various ssDNA-binding proteins (SPATA22, MEIOB and RPA) in HSF2BP and BRME1 immunoprecipitates further supported the model, and led to the suggestion that HSF2BP and BRME1 act as adaptors, anchoring BRCA2 to the protected ssDNA. Such adaptors have not been identified in somatic HR. Both BRCA2 and its equally important partner PALB2, also involved in meiotic HR (Hartford et al., 2016), are well equipped for the recruitment to resected DSBs with a set of DNA-, RAD51- and chromatin-binding domains, and can stimulate RAD51 and/or DMC1 activity autonomously *in vitro*. As a localizer, HSF2BP neither mimics DNA as does the BRCA2-interacting protein DSS1 (Zhao et al., 2015), nor interacts with ssDNA or dsDNA (**Supplementary Figure S9**; see also (Felipe-Medina et al., 2020)). However, even with these reservations regarding the localizer model, we fully expected the HSF2BP-binding domain of BRCA2 to be essential for meiotic HR, because it is conserved, and no somatic function could be assigned to it by several previous studies in human cancer cells (Li et al., 2009; Tavtigian et al., 2008). The high evolutionary conservation of the BRCA2-HSF2BP interaction, initially based on sequence analysis and interchangeability of human and frog HSF2BP proteins in biochemical experiments, is further emphasized by the structure we solved, which revealed that the HSF2BP surface binding to BRCA2 is also highly conserved (**Figure 3C**). Further analysis of the function of HSF2BP and the role of the HSF2BP-BRCA2 interaction in meiosis, and outside of it, is required to explain these conservations.

The crystal structure of the complex revealed the large size of the interface between BRCA2 and HSF2BP, and we consistently measured a nanomolar affinity between the two proteins. However, analysis of low frequency human polymorphisms revealed that single amino acid substitutions in BRCA2 and HSF2BP can be sufficient to disrupt the interaction, which might be clinically relevant for patients with either fertility defects or cancers. Finally, most remarkably, our structural and biochemical data revealed that binding triggers oligomerization of HSF2BP. It was already reported that full length HSF2BP contains an N-terminal domain that dimerizes (Zhang et al., 2020). Here, we identified two additional mechanisms of oligomerization: the ARM domain of HSF2BP is dimeric, and it tetramerizes upon BRCA2 binding. Thus, multiple oligomeric structures of HSF2BP can result from its binding to BRCA2. Moreover, at least four HSF2BP molecules bind to two BRCA2, so that one HSF2BP oligomer clearly binds to more than one BRCA2 protein. The interaction triggers HSF2BP oligomerization, but this oligomer includes several BRCA2 molecules, which might favor full length BRCA2 oligomerization as well. Existence of three distinct oligomerization mechanisms, each of which can be under separate regulatory control, allows HSF2BP to serve as a versatile and potent agent increasing the local concentration and/or modifying the oligomerization state of BRCA2. This can play a positive role in some contexts, and be detrimental in others. For example, we previously found that HSF2BP, when produced ectopically in somatic cancer cells, interferes with the role of BRCA2 in DNA interstrand crosslink repair by causing its degradation (Sato et al., 2020). The findings we report here suggest that this degradation, which is mediated by the p97 segregase and is proteasome-dependent, could result from HSF2BP-induced BRCA2 aggregation.

Altogether we conclude that the evolutionarily conserved high-affinity oligomerization-inducing interaction between HSF2BP and BRCA2 we described in this paper is not required for the recruitment of RAD51 and DMC1 strand exchange proteins and for productive HR in meiosis. The meiotic function of HSF2BP that is associated with fertility may therefore result from other interactions, mediated either by its N-terminal coiled-coil domain, which binds BRME1, by the conserved surface of its ARM domain binding to other proteins with sequences similar to the cryptic repeat our structure revealed in BRCA2, or by other parts of the ARM domain.

## Acknowledgments

We thank Ambre Petitalot for the first purification of the BRCA2 F0 fragment, Christophe Velours on the I2BC protein-protein interaction facility for the SEC-MALS experiments, Guillaume Hoffmann and Jose A. Marquez for their advices during crystallogenesis assays on the HTX platform, and Virginie Ropars for her great help during crystallogenesis assays, crystal freezing and X-ray crystallography data collection. We acknowledge SOLEIL for provision of synchrotron radiation facilities and we would like to thank the respective staffs for assistance in using PROXIMA-1, PROXIMA-2 and SWING beamlines. We thank Nicole van Vliet for the assistance with the initial mouse experiments.

## Materials and Methods

### Cells, DNA Constructs and Transfection

HeLa (human cervical adenocarcinoma, female origin) and HEK293T (human embryonic kidney, female origin) cells were cultured in DMEM supplemented with 10% FCS, 200 U/ml penicillin, 200 μg/ml streptomycin. mES cell lines were derived from the IB10 cell line, which is a subclone of E14 129/Ola from male origin (Hooper et al., 1987), specific pathogen free. Cells were cultured on gelatinized plastic dishes (0.1% gelatin in water) as described before (Zelensky et al., 2017) at atmospheric oxygen concentration in media comprising 1:1 mixture of DMEM (Lonza BioWhittaker Cat. BE12-604F/U1, with Ultraglutamine 1, 4.5 g/l Glucose) and BRL-conditioned DMEM, supplemented with 1000 U/ml leukemia inhibitory factor, 10% FCS, 1x NEAA, 200 U/ml penicillin, 200 μg/ml streptomycin, 89 μM β-mercaptoethanol.

Expression constructs for producing point mutation and truncation variants of HS2BP and BRCA2 in human cells were engineered as described before (Brandsma et al., 2019) in the PiggyBac vectors by Gibson assembly. For transient expression into HEK293T cells plasmid DNA was transfected using calcium precipitation method or PEI transfection. For stable integration into HeLa cells, PiggyBac expression vectors were co-lipofected together with PiggyBac transposase plasmid (hyPBase, (Yusa et al., 2011)) with Lipofectamine 3000 (Thermo Fisher).

GFP knock-in alleles in mES cells were engineered using the previously described CRISPR/Cas9-stimulated approach. All gRNAs were cloned into a derivative of pX459 vector. Excision of BRCA2 exon 12 from mES cells was performed with gRNAs targeting the same sequences in intron 11 and intron 12 as those used to produce *Brica2*Δ12 mouse alleles (see below). Excision of exons 12-14 was performed with gRNAs targeting the same sequence in intron 11, and the sequence in intron 14 CCAACCAGCCCGGTCAAGTT. IB10 or the previously described *Brca2^GFP/GFP^* (Reuter et al., 2014) were used as parental cell lines. Excision of exons 12-14 from HeLa cells was performed with the same gRNA target in intron 11 as used before and the following target in intron 14: AGGAGAGCATGTAAACTTCG. Cells were genotyped by PCR. Excision was further confirmed by RT-PCR analysis of the first-strand cDNA produced from total mRNA with oligo-dT primers with SuperScript II polymerase (Invitrogen).

### Immunoprecipitation and immunoblotting

Cells were washed twice in ice-cold PBS and lysed in situ in NETT buffer (100 mM NaCl, 50 mM Tris pH 7.5, 5 mM EDTA pH 8.0, 0.5% Triton-X100) supplemented immediately before use with protease inhibitor cocktail (Roche) and 0.4 mg/ml Pefabloc (Roche) (NETT++); 450 fifty μl NETT++ buffer was used per 145 mm dish ES cells, 1 ml for HeLa and HEK293T. After 5-10 min, cells were scraped off and collected in 1.5 microcentrifuge tubes; lysis was continued for additional 20-30 min on ice, then mixtures were centrifuged (15 min, 4º, 14000 rcf) and the supernatant (input) was added to washed anti-GFP beads (Chromotek). Beads and lysates were incubated 2-4 h at 4 °C while rotating, washed three times in NETT++ buffer and bound proteins were eluted by boiling in 2x Sample buffer. Immunoblotting was performed following standard procedures with the following antibodies: anti-GFP mAb (Roche, #11814460001), anti-GFP pAb (Abcam #ab290 and Invitrogen #A11122) anti-RAD51 (Essers et al., 2002)anti-BRCA2 mAb Ab1 OP-95 (Millipore #OP95), anti-BRCA2 (Abcam #27976), anti-HSF2BP (Brandsma et al., 2019), anti-Flag (M2 antibody, Sigma, F3165). For quantitative immunoblotting fluorescently labeled secondary antibodies were used: anti-mouse CF680 (Sigma #SAB460199), anti-rabbit CF770 (Sigma #SAB460215); membranes were scanned using Odyssey CLx imaging system (LI-COR).

### Protein expression and purification

Human full length HSF2BP WT and R200T were expressed using a pETM11 (6xHis-TEV) expression vector in *E. coli* BL21 DE3 Rosetta2 cells, and purified as previously reported (Brandsma et al., *Cell Rep* 2019). The armadillo domain of HSF2BP, from aa 122 to aa 334, which we will further name ARM, was similarly expressed using a pETM11 (6xHis-TEV) expression vector in BL21 DE3 Rosetta2 cells, and purified as full length HSF2BP.

Human BRCA2 fragment F0 was expressed using a pET-22b expression vector as a fusion protein comprising BRCA2 from aa 2213 to aa 2342 (including mutation C2332T to avoid oxidation problems), a TEV site, GB1 and 6xHis, in *E. coli* BL21 DE3 Star cells. The BRCA2 gene was optimized for expression in bacteria and synthetized by Genscript. In addition, smaller BRCA2 fragments were produced using the same strategy: F_NMR_ and F15X corresponding to aa 2252 to aa 2342 and aa 2291 to aa 2343, respectively. The smallest fragment F15XΔ12, from aa 2312 to aa 2342, was synthetized by Genecust.

For NMR analysis, ^15^N-labeled or ^15^N- and ^13^C-labeled BRCA2 fragments were produced in *E. coli* BL21 DE3 Star cells grown in M9 medium containing either 0.5 g/l ^15^NH_4_Cl or 0.5 g/l ^15^NH_4_Cl and 2 g/l ^13^C-glucose, respectively. For crystallography, selenomethionine(SeMet)-labeled ARM and F15X were produced in transformed *E. coli* BL21(DE3) Rosetta2 (ARM) and Star (F15X) cells, Respectively, grown in minimum medium (16 g of Na_2_HPO_4_, 4 g of KH_2_PO_4_, 1 g of NaCl, 0.5g of EDTA, 0.4 g of FeCl_3_, 0.04 g of ZnCl_2_, 0.006 g of CuCl_2_ 6 H_2_O, 0.005 g of CoCl_2_, 0.005 g of H3BO3, 4 g of glucose, 20 mg of thiamine, 20 mg of biotin, 1g of (NH_4_)2SO_4_, 0.5 g of MgSO_4_, and 0.1 g of CaCl_2_ in 1 liter of MilliQ), supplemented with 200 mg of each amino acid and 125 mg of SeMet per 1 liter of medium.

Protein expression was induced with 0.2 mM IPTG at an OD_600_ of 0.6, and incubated further ON at 20°C (HSF2BP), or induced with 1 mM IPTG and incubated for 3h at 37°C (BRCA2). Harvested cells were resuspended in 25 mM Tris-HCl pH 7.5.,500 mM NaCl (HSF2BP) or 150 mM NaCl (BRCA2), 5 mM β-mercaptoethanol, EDTA-free Protease Inhibitor Cocktail (Roche) and disrupted by sonication. Lysates were supplemented with 1.5 mM MgCl_2_ and treated by Benzonase nuclease at 4°C for 30 min, and then centrifuged at 20.000 rpm at 4 °C for 30 min. After filtration (0.4 μm), the supernatant was loaded on a chromatography HisTrap HP 5 mL column (GE Healthcare) equilibrated with the buffer Tris-HCl 25 mM. pH 8, 500 mM NaCl (HSF2BP), or 150 mM NaCl (BRCA2) and 5 mM β-mercaptoethanol. The proteins were eluted with a linear gradient of imidazole. The tag was cleaved by the TEV protease (at a ratio of 2 % w/w) during an ON dialysis at 4°C against 25 mM Tris-HCl pH 7.5,150 mM NaCl, 5 mM β-mercaptoethanol. The protein solution was loaded on a HisTrap column and the tag free proteins were collected in the flow through. Finally, a size exclusion chromatography was performed on HiLoad Superdex 10/300 200pg (HSF2BP) or 75pg (BRCA2) equilibrated in 25 mM Tris-HCl pH 7.5, 250 mM NaCl (HSF2BP) or 150 mM NaCl (BRCA2), 5 mM β-mercaptoethanol. The quality of the purified protein was analysed by SDS-PAGE and the protein concentration was determined by spectrophotometry using the absorbance at 280 nm. The protein thermal stability was evaluated using the simplified Thermofluor assay available on the High Throughput Crystallisation Laboratory (HTX Lab) of the EMBL Grenoble (Dupeux et al., 2011).

### BRCA2 peptide structural analysis by NMR

The ^15^N/^13^C labeled F0 fragment was analyzed by 3D heteronuclear NMR. in order to assign its Hn, N, Cα, Cβ and Co chemical shifts, and identify its binding site to HSF2BP. Therefore, 3D NMR HNCA, CBCACONH, HNCACB, HNCANNH, HNCO and HNCACO experiments were performed on a 700 MHz Bruker AVANCE NEO spectrometer equipped with a triple resonance cryogenic TCI probe at 283 K. The data were processed using Topspin 4.0 (Bruker) and analyzed using CCPNMR 2.4 (Vranken et al., *Proteins* 2005). Sodium trimethyl-silyl-propane-sulfonate (DSS) was used as a chemical shift reference. Experiments were performed on a 5-mm-diameter Shigemi sample tube containing the 500 μM uniformly ^15^N/^13^C-labeled protein, in 40 mM sodium phosphate pH 7.0, 150 mM NaCl and 95:5% H_2_O/D_2_O. For binding studies, 2D NMR ^1^H-^15^N HSQC experiments were recorded on a 3-mm-diameter sample tube containing the ^15^N-labeled F0 at 100 μM in the absence and presence of the ARM domain at 100 μM, on a 950 MHz Bruker Avance III spectrometer equipped with a triple resonance cryogenic TCI probe at 283 K.

### Protein-protein interactions

Isothermal Titration Calorimetry (ITC) experiments were performed using a high-precision VP-ITC instrument (GE Healthcare) at 293 K. To characterize the interactions between HSF2BP and BRCA2 fragments (WT, variants), the proteins were dialyzed against 25 mM Tris pH 7.5, 250 mM NaCl, 5mM β-mercaptoethanol. The HSF2BP (or ARM domain) was in the sample cell at 10-20 μM and was titrated by BRCA2 fragments at 40-100 μM, using 10 μL injections with 210 s intervals between each injection. The first 2 μL injection was ignored in the final data analysis. The integration of the peaks corresponding to each injection. the correction for the baseline and the fit were performed using the Origin 7.0 software provided by the manufacturer, to obtain the stoichiometry (N), dissociation constant (K_d_) and enthalpy of complex formation (ΔH) for each interaction. These data are indicated in Table 1. Two replicates were performed for each experiment.

Size-Exclusion Chromatography (SEC) coupled to Multi Angle Light Scattering (MALS) was used in order to measure the molecular masses of the complexes in solution. Therefore, HSF2BP and ARM proteins were loaded in the presence and absence of the BRCA2 fragment F15X on a Superdex 200 10/300 GL (GE Healthcare) using a HPLC Shimadzu system coupled to MALS/QELS/UV/RI (Wyatt Technology). The chromatography buffer was 25 mM Tris-HCl buffer, pH 7.5, 250 mM NaCl, 5mM β-mercaptoethanol. The proteins were injected at 0.8 to 1 mg/ml in 100 μl. Data were analyzed using the ASTRA software; a calibration was performed with BSA as a standard.

Size-Exclusion Chromatography (SEC) coupled to Small Angle X-ray Scattering (SAXS) is available on the SWING beamline at synchrotron SOLEIL, in order to obtain a distance distribution corresponding to each sample in solution. The free HSF2BP protein as well as the complex between ARM and F15X were analyzed using a Bio SEC-3 column (Agilent) equilibrated in 25 mM Tris-HCl buffer, pH 7.5, 250 mM NaCl, 5 mM β-mercaptoethanol. The proteins were loaded at a concentration of 6 and 10 mg/ml, in order to observe an elution peak at an OD_280nm_ of 1 and 1.2 AU, respectively.

Size-Exclusion Chromatography (SEC) was also used to characterize ARM and ARM bound to F15XΔ12. The ARM domain was loaded on a Superose 6 Increase 10/300 (GE Healthcare) alone (at 2 mg/ml) or bound to F15XΔ12 (at 5 mg/ml, ARM to peptide ratio 1:1.2) in 25 mM Tris-HCl buffer, pH 7.5, 250 mM NaCl, 5mM β-mercaptoethanol. The OD values from the elution of ARM alone were multiplied by 2.5 to be compared to the OD values from the elution of ARM bound to F15XΔ12.

### Crystallization and structure determination

Prior to crystallization, the complex between ARM and F15X was loaded on a size exclusion chromatography column HiLoad Superdex 16/600 200pg (GE Healthcare) equilibrated in a 25 mM Tris pH 7.5, 250 mM NaCl and 5mM β-mercaptoethanol, in order to prevent the presence of aggregates. The complex was then concentrated up to 10 mg/ml. Initial crystallization experiments were carried out at the High Throughput Crystallisation Laboratory (HTX Lab) of the EMBL Grenoble (Dimasi et al., 2007). Crystals were prepared for X-ray diffraction experiments using the CrystalDirect technology (Zander et al., 2016). Crystals were obtained using the hanging-drop vapor diffusion method at 291 K. One μl of protein and reservoir solution containing 100 mM MgCl_2_, 100 mM MES pH 6 and 16% (w/v) PEG 3350 were mixed. Needle crystals appeared within 3 days, were grown for 1–2 weeks and were frozen in liquid nitrogen after cryoprotection using the reservoir solution supplemented with 20% glycerol.

X-ray diffraction data were collected on the beamlines PROXIMA-1 and PROXIMA-2 at the SOLEIL synchrotron (St Aubin, France) and reduced using the XDS package (Kabsch, 2010). First phases for a triclinic crystal form (Table 2) were obtained by molecular replacement using the program MOLREP from CCP4 (Winn et al., 2011) and different homologous models. One of the models obtained by the Robetta server (Song et al., 2013) gave the best correlation in the final translation function. These phases allowed to find the selenium substructure from a SeMet SAD dataset from the same crystal form with only ARM protein containing selenomethionines. Later a monoclinic crystal form (Table 2) was obtained from a complex with both ARM and F15X containing selenomethionines. The collected SeMet SAD dataset (wavelength of data collection: lambda=0.97918 Å) allowed to directly calculate phases, without external model contributions, and confirmed the initial model built in the triclinic crystal form. Se sites were found using the SHELX C/D/E suite of programs. These sites were refined using PHASER in EP mode. The resulting Se SAD phases were improved by density modification using PARROT and a model automatically build using BUCCANEER confirming the sequence attribution for ARM and F15X. The resulting model underwent iterative cycles of manual reconstruction in COOT (Emsley et al., 2010) and refinement in BUSTER ((Bricogne et al., 2020); Table 2). At the end of the refinement, 90% and 3.5% of the residues were in favored and outlier regions of the Ramachandran plot, respectively. Few residues were not visible in the electron density (L55 in chain B, V52 and Ala53 in chain C, loop 51-55, F169, and R208 in chain D, and H33 in chain F). These residues were included in the pdb file, but with an occupancy of 0. The final pdb file and monoclinic data set have been deposited in the Protein Data Bank (entry code 7BDX).

### Meiotic spread nuclei preparations and immunocytochemistry

Meiotic testicular cells were spread as previously described (Peters et al., 1997). For immunocytochemistry the slides were washed in PBS (3x 10 min), blocked in 0.5% w/v BSA and 0.5% w/v milk powder in PBS followed by staining with primary antibody which was diluted in 10% w/v BSA in PBS and incubated overnight at room temperature in a humid chamber. Subsequently, the slides were washed with PBS (3x 10 min), blocked in 10% v/v normal goat serum (Sigma) in blocking buffer (supernatant of 5% w/v milk powder in PBS centrifuged at maximum speed for 10 min) followed by staining with secondary antibody which was diluted in 10% v/v normal goat serum (Sigma) in blocking buffer and incubated for 2 hours at room temperature in a humid chamber. Finally, the slides were washed with PBS (3x 10min) and embedded in Prolong Gold with DAPI (Invitrogen). The following primary antibodies were used: anti-DMC1 (1:1000,Abcam ab11054), mouse anti-SYCP3 (1:200, Abcam ab97672), mouse anti-MLH1 (1:25, BD Pharmingen 51-1327GR), and rabbit polyclonal anti-RAD51 (1:1000, (Essers et al., 2002)), rabbit anti-HSF2BP (1:30, #1, (Felipe-Medina et al., 2020)) and rabbit anti-BMRE1 (1:100, #2, (Felipe-Medina et al., 2020)), rabbit polyclonal anti-SYCP3(1:5000, (Lammers et al., 1994)) and guinea pig anti-SYCP2 (1:100, (Winkel et al., 2009)) and guinea pig anti-HORMAD2 (1:100, (Wojtasz et al., 2009)). Secondary antibodies: goat anti-guinea pig Alexa 546, goat anti-rabbit Alexa 488, 555 and goat anti-mouse Alexa 488, 555 and 633.

Immunostained spreads were imaged using a Zeiss Confocal Laser Scanning Microscope 700 with 63x objective immersed in oil. All images within one analysis were taken with the same intensity. Images were analyzed using ImageJ (Fiji) software. RAD51 and DMC1 foci quantification was performed using the ImageJ function “Analyze particles” in combination with a manual threshold. MLH1 foci were counted manually. BMRE1 foci quantification was performed using the ImageJ function “Find Maxima” with a prominence of 40, and a mask of SYCP3 was used to reduce the background signal. Also for the HSF2BP intensity quantification, a mask of SYCP3 was used and the mean HSF2BP intensity within this mask was measured.

### Animals

All animals were kept in accordance with local regulations under the work protocols 17-867-11 and 15-247-20. Animal experiments were approved by the Dutch competent authority (Centrale Commissie Dierproeven, CCD) and all experiments conform to relevant regulatory standards. Female mice for CRISPR/Cas9 injection were C57BL/6 OlaHsd from Envigo, age 5 weeks. For spermatogenesis analysis male mice were sacrificed at the age of 15 weeks.

### Brca2-Δ12 mouse generation

*Brca2* Δ12 mice were generated by two CRISPR/Cas9 cut excision, as described before (Brandsma et al., 2019). Female donor mice (age 5 weeks, C57BL/6 OlaHsd from Envigo) were superovulated by injecting 5-7.5 IE folligonan (100-150 μl, IP (FSH hormone; time of injection ± 13.30 h; day −3). Followed at day −1 by an injection of 5-7.5 IE chorulon (100-150 μl, IP (hCG hormone; time of injection 12.00 h). Immediately after the chorulon injection, the females were put with fertile males in a one to one ratio. Next day (0) females were euthanized by cervical dislocation. Oviducts were isolated, oocytes collected and injected with ribonucleoprotein complexes of S.p.Cas9 3NLS (IDT cat. no. 1074181), crRNA and tracrRNA (both Alt-R, synthesized by IDT). Target sequences for crRNA were TAATATTCCAACCCTCGTGT (upstream of *Brca2* exon 12) and TGAGAAATGTACACCTCATT (downstream of exon 12). For ribonucleoprotein formation equal volumes (5 μL) of crRNA and tracrRNA (both 100 μM in IDT annealing buffer) were mixed, heated to 95 ºC for5 min and allowed to cool on the bench. The annealed RNAs (1.2 μL, 50 μM) were mixed with Cas9 (10 μl diluted to 200 ng/μl in the DNA microinjection buffer (10 mM Tris-HCl, pH 7.4, 0.25 mM EDTA in water) at the final concentrations 0.12 μM Cas9, 0.6 μM of each of the two crRNA:tracRNA complexes in microinjection buffer. Foster mother (minimum age 8 weeks) were set up with vasectomized males in a 2 to 1 ratio. Next day (0), pseudopregnant female (recognized by a copulation prop) were collected. For transplanting the injected oocytes, pseudopregnant female was anesthetized by an IP injection of a mix of Ketalin (12 mg/ml ketamine in PBS)-Rompun (0.61 xylazine mg/ml PBS) 100 μl per 10 g bodyweight). Post-surgery pain relief was given when the mouse was anaesthetized (S.C. Rimadyl Cattle, 5mg/ml in PBS, dose 5μg/g mouse). Transplantation resulted in 8 pups from a single litters, of which 3 (all female) contained the deletions in the targeted region, as determined by PCR genotyping. Different primer combinations were used for initial genotyping, but mB2i11-F1 AGCTGCCACATGGATTCTGAG, mB2i12-R2 GGACTAAGAGGCAAGGCATCA, and mB2e12-R1 GCTTTTTGAAGGTGTTAAGGATTTT, were used routinely. Sequencing of the PCR products from the founder animals revealed mosaicism for the junctions between the two CRISPR/Cas9 cuts; deletion sizes were close to expected 713bp, bigger (1179 bp) or smaller due to insertions of ectopic DNA. The experimental cohort was eventually formed through back-crossing and inter-crossing from one founder, with the deletion produced by direct ligation between the two Cas9 cuts. Routine PCR genotyping of was performed using MyTaq Red mix (Bioline) and using the mentioned primers in 1:0.5:1 combination, for simultaneous amplification of the wild-type and the Δ12 alleles (PCR products 663 and 314 bp, respectively).

Adult wild type and *Brca2^Δ12^* males were sacrificed and weighed, and testes and epididymides were collected and also weighed. Epididymides were collected in PBS, dounced, and sperm cells were counted. For fertility assessment breedings were set up between *Brca2^Δ12/Δ12^* and wild-type C57BL/6 animals.

### Immunofluorescence and Microscopy

Immunofluorescence staining was performed on ES cells grown overnight on a glass coverslip coated with laminin. Sterile 24 mm coverslip was placed in a 6-well plate, and a 100 μl drop of 0.05 mg/ml solution of laminin (Roche, 11243217001) was pipetted in the middle of it. The plate was left for ^~^30 min in the cell culture incubator, after which the laminin solution was aspirated, and cell suspension was placed in the well. DNA damage was induced by irradiation with 8 Gy X-ray followed by 2 h recovery. Cells were washed with PBS, pre-extracted in sucrose buffer (0.5% Triton X-100, 20 mM HEPES pH 7.9, 50 mM NaCl, 3 mM MgCl_2_, 3 mM sucrose) for 1 min, fixed for 15 min in 2% paraformaldehyde in PBS at room temperature, immunostained with anti-RAD51antibody and mounted with DAPI. Images were acquired using Leica SP8 confocal microscope in automatic tile scan mode. Maximum projections from a z-stack of 3 confocal planes through a 1 μm slice were produced for analysis. HSF2BP-GFP foci were quantified automatically using CellProfiler (Carpenter et al., 2006). In short, nuclei were segmented using a global threshold (minimum cross-entropy) based on the DAPI signal. The masked images were used to identify HSFP2BP foci using a global threshold (Robust background) method with 2 standard deviations above background. Subsequently the number of foci was counted per segmented nucleus.

For single particle tracking, cells were grown overnight in 8-well glass bottom dishes (Ibidi) coated with 0.05 mg/ml laminin. Prior to the experiment cell medium was replaced with imaging medium (Fluorobrite DMEM (Thermo Fisher), complemented with 10% FCS, 1x NEAA, 89 μM β-mercaptoethanol, 200 U/ml penicillin, 200 μg/ml streptomycin, and 1,000 U/ml leukemia inhibitory factor). Live-cell experiment was performed on a Zeiss Elyra PS complemented with a temperature-controlled stage and objective heating (TokaiHit). Samples were kept at 37 ºC and 5% CO_2_ while imaging. For excitation of GFP a 100mW 488 nm laser was used. The samples were illuminated with HiLo illumination by using a 100x 1.57NA Korr αPlan Apochromat (Zeiss) TIRF objective. Andor iXon DU897 was used for detection of the fluorescence signal, from the chip a region of 256 by 256 pixels (with an effective pixel size of 100*100 nm) was recorded at 19.2 Hz interval (50 ms integration time plus 2 ms image transfer time). EMCCD gain was set at 300. Per cell a total of 200 frames were recorded.

## Supplementary Figure Legends

**Supplementary Figure S1.**
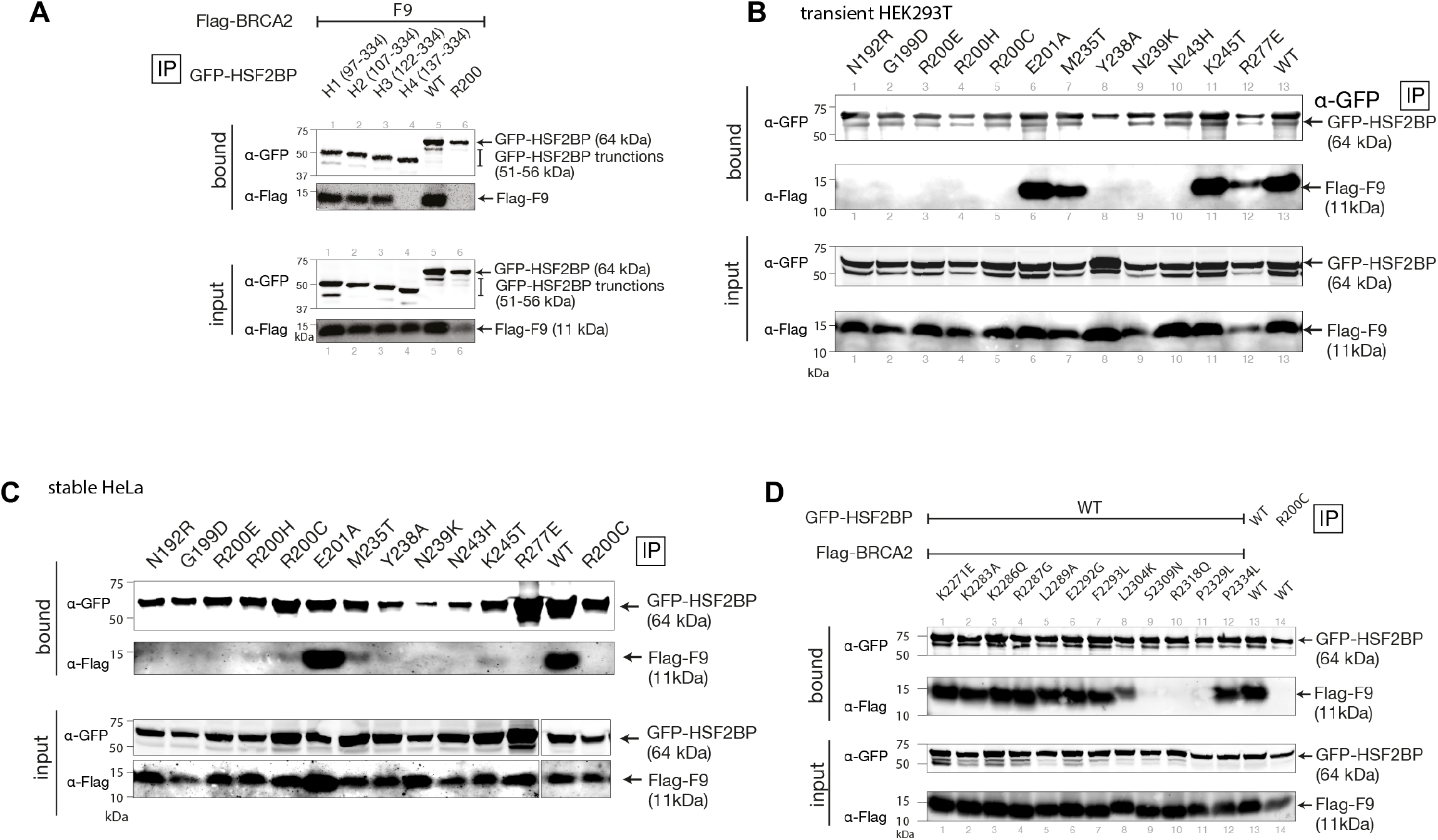
Co-immunoprecipitation of HSF2BP and BRCA2 variants. **(A)** Full-length GFP-HSF2BP and its truncated versions (H1-H_4_) and Flag-tagged BRCA2 fragment F9 were transiently overproduced in HEK293T cells, co-immunoprecipitated using anti-GFP beads and analyzed by immunoblotting with anti-GFP and anti-Flag antibodies. **(B)** Co-immunoprecipitation experiment from HEK293T cell lines transiently overproducing GFP-HSF2BP point mutants and a Flag-BRCA2 fragment corresponding to the previously identified minimal HSF2BP-binding domain (HBD, G2270-T2337. (Brandsma et al., 2019)). (**B**) As in panel (A), but fragments were produced by stable transformation of HeLa cells. **(C)** Co-immunoprecipitation experiment from HEK293T cell lines transiently overproducing GFP-HSF2BP and variants of the Flag-BRCA2-F9 fragment.

**Supplementary Figure S2.**
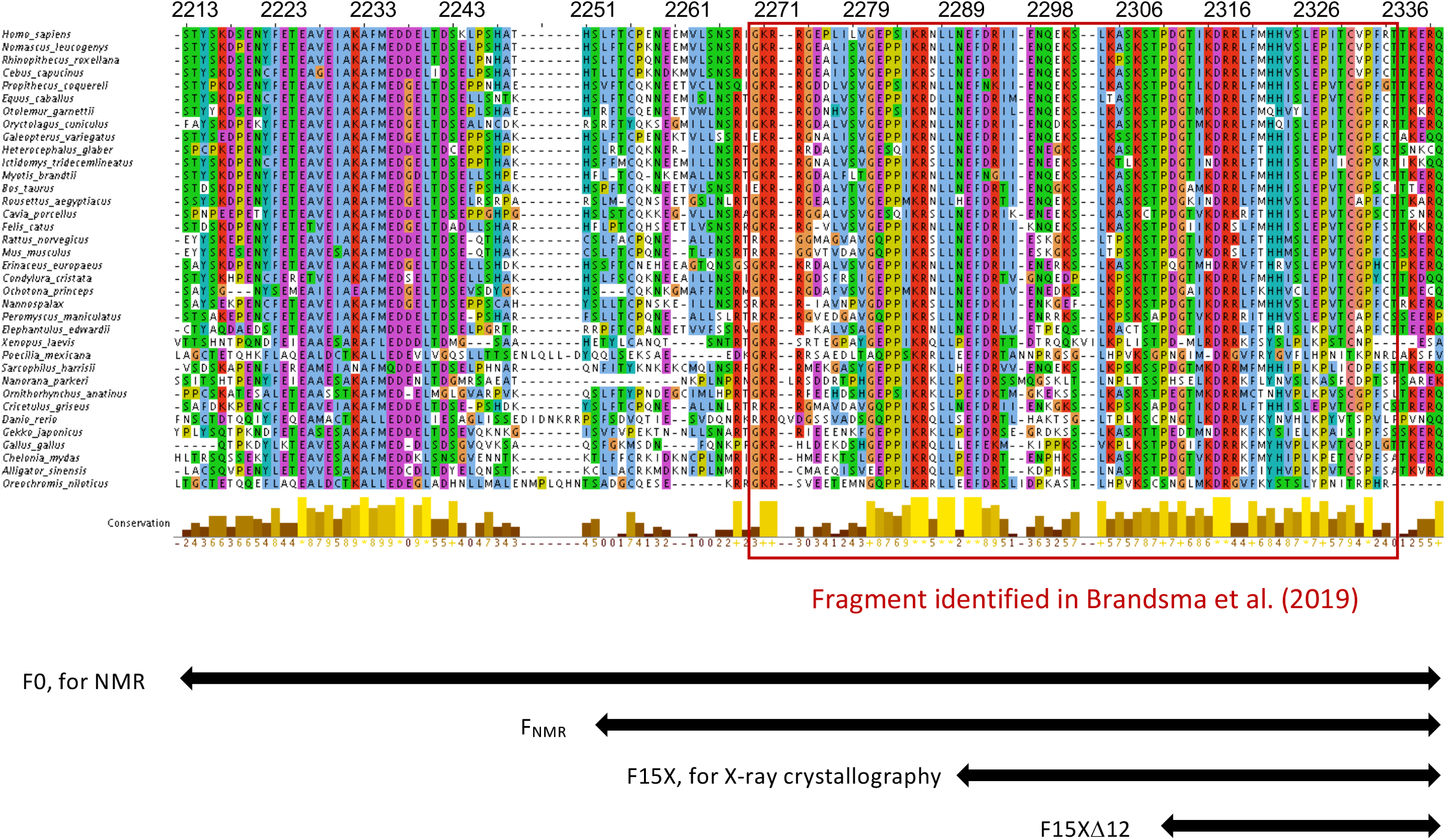
Sequence alignment of 36 BRCA2 homologs focused on region F0 (S2213-Q2342) of human BRCA2. Alignment and scoring were calculated using Jalview 2.10.1. The fragment previously identified as responsible for HSF2BP binding is boxed. The fragments produced for this study are indicated below.

**Supplementary Figure S3.**
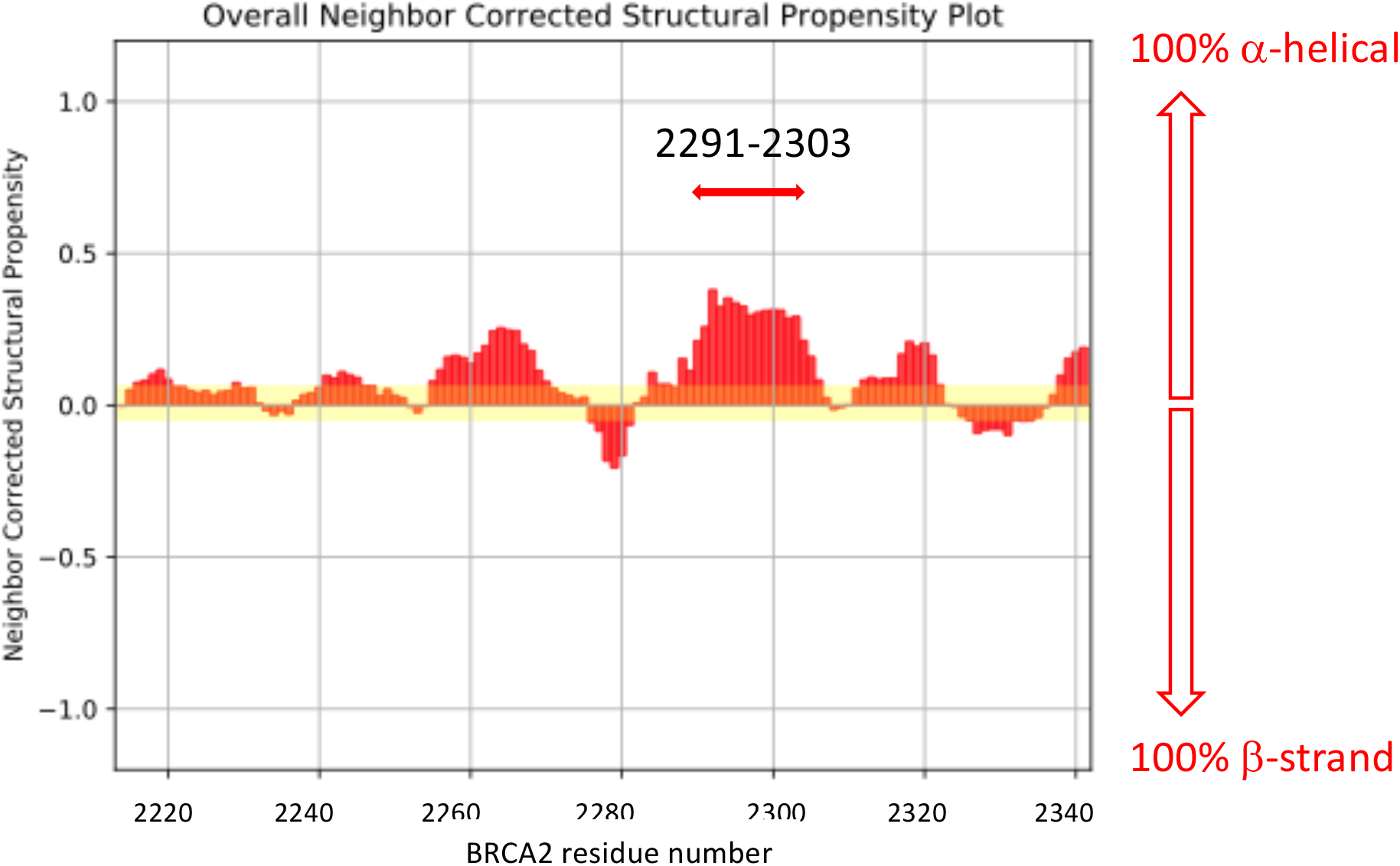
BRCA2 fragment F0 is disordered in solution. Analysis of the 3D heteronuclear NMR experiments recorded on the ^15^N, ^13^C labeled BRCA2 fragment F0 (S2213-Q2342) provided the chemical shifts for all Hn, N, Ca, Cb and Co nuclei. From these chemical shifts, the neighbor corrected structural propensity was calculated as a function of BRCA2 residue number, using the Webserver called “neighbor corrected structural propensity calculator” (https://st-protein02.chem.au.dk/ncSPC/; (Tamiola et al., 2010)). Positive and negative values correspond to populations with α-helical and β-strand conformations, respectively. In the case of BRCA2 F0, propensity values are all lower than 0.5, indicating that the protein is disordered in solution. Only region N2291-S2303 forms a transient α-helix present in more than 25% of the molecules.

**Supplementary Figure S4.**
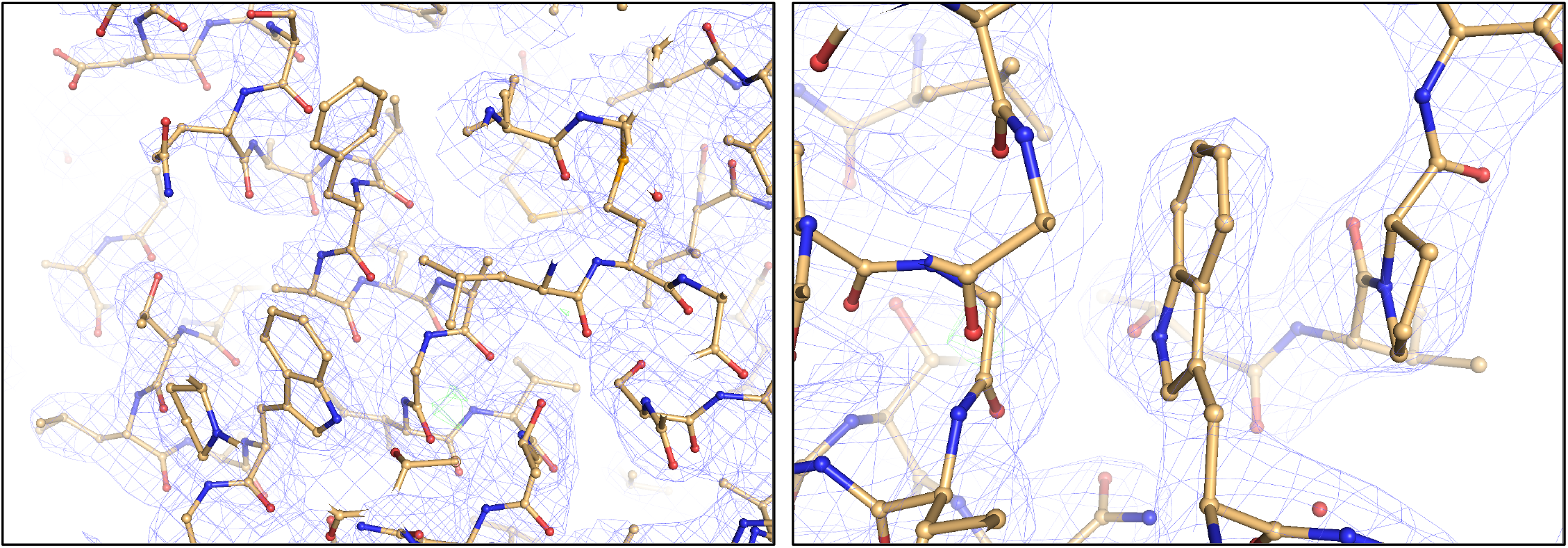
The crystal structure is nicely defined by the 2.6 Å resolution electron density. Example of the quality of the 2FoFc electron density map at 1.0 RMSD, showing a stacking between a tryptophane and a proline.

**Supplementary Figure S5.**
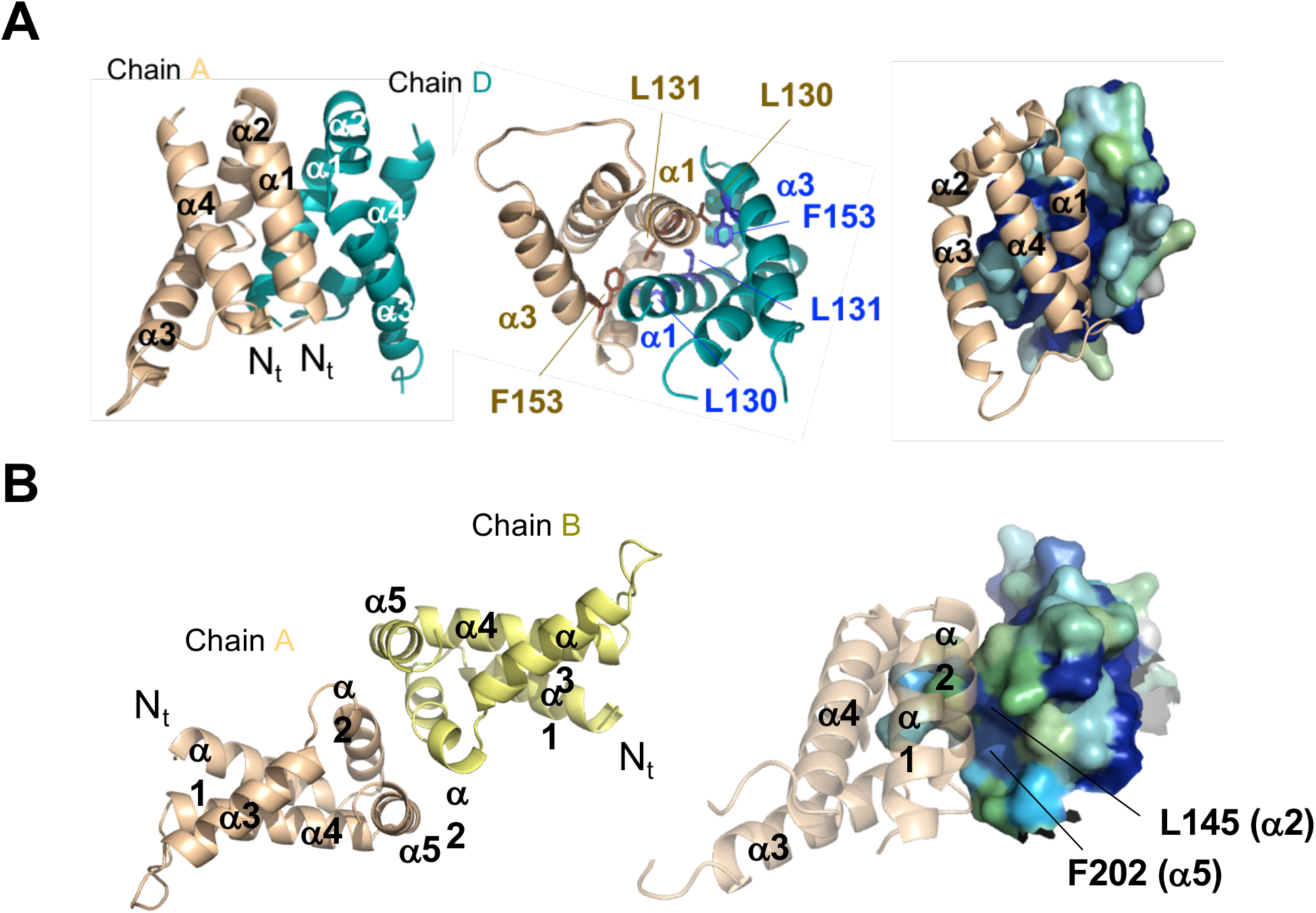
Oligomerization interfaces within the complex. **(A)** Dimerization interface observed between chains A and D. A similar interface is observed between chain B and C. In the left panel, the four N-terminal α-helices involved in the interface are displayed. In the middle panel, the three conserved residues of the interface are shown in sticks. In the right panel, the surface of chain D N-terminal helices is colored as a function of scores calculated by Consurf (Ashkenazy et al., NAR 2016). High, medium, weak and no conservation are indicated using dark blue, cyan, green and grey, respectively. **(B)** Central interface observed between chains A and B, resulting from the tetramerization of the ARM domains. Left panel: the symmetric interface of about 250 Å^2^ involves helices α2 (K142, A143, K149 and A150) and helices α5 (N205, S206). Right panel: analysis of this interface using Consurf showed that it mostly comprises poorly conserved residues (see the surface representation of chain B). Only L145 in helix α2 and F202 in helix α5 are conserved; they interact with each other within each ARM chain, thus contributing to the interface between α2 and α5; however, they do not interact with conserved residues from the other chain.

**Supplementary Figure S6.**
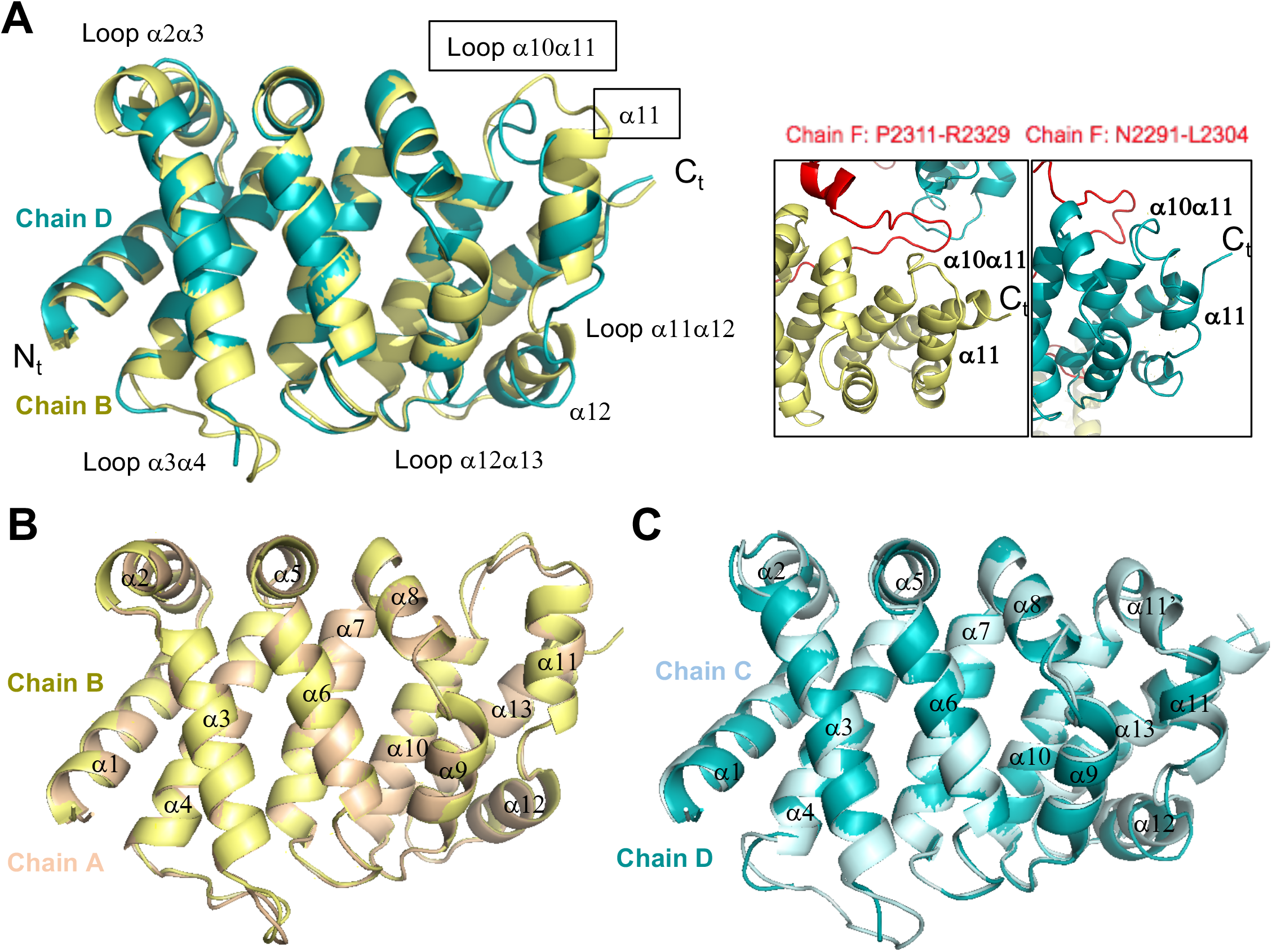
Structural variations are observed between ARM chains interacting with the N-terminal region of BRCA2 F15X and ARM chains interacting with the C-terminal region of BRCA2 F15X. (**A**) Superimposition of the structures of chains B and D, interacting with different parts of the BRCA2 peptide. Variations are observed, in particular in loop α10α11 and helix α11. These structural elements interact with either BRCA2 fragment P2311-R2319, or BRCA2 fragment N2291-L2304, as indicated in the boxed view. (**B, C**) Superimposition of the structures of (B) chains A and B, and (C) chains C and D, interacting with similar parts of the BRCA2 peptide. These structures are highly similar, as demonstrated by the low RMSD values measured between their backbone Cα atoms, displayed in Figure 3A.

**Supplementary Figure S7.**
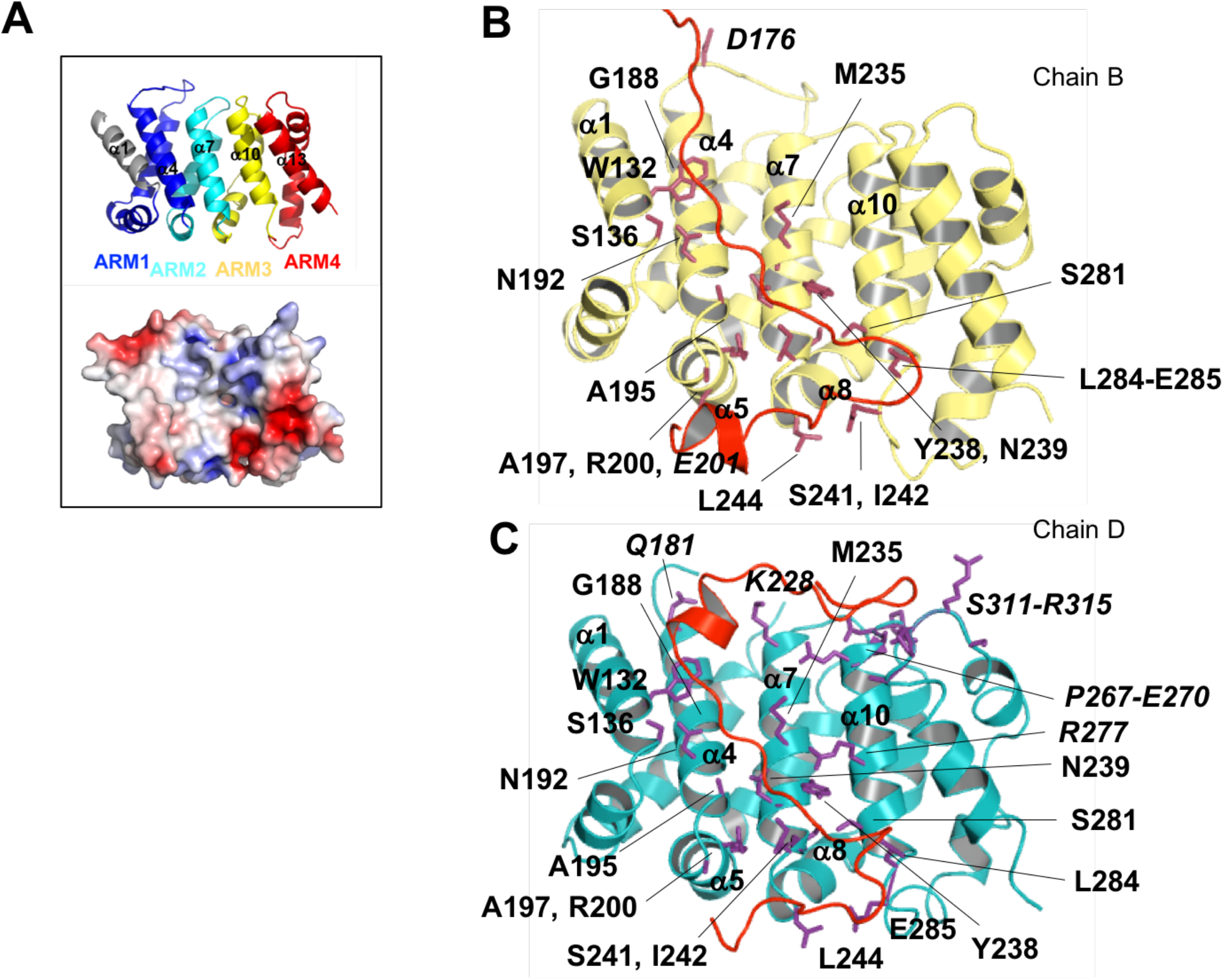
A set of ARM residues interact with both the N-terminal and C-terminal regions of the BRCA2 peptide. **(A)** Cartoon view of the four Armadillo repeats of ARM, colored in blue, cyan, yellow and red. The electrostatic potential at the surface of the domain is displayed at the bottom of the panel, to highlight the positively charged character of the groove defined by helices α1, α4, α7, α10 and α13. **(B,C)** Zoom on the interfaces between **(B)** chain B and the C-terminal region of F and **(C)** chain D and the N-terminal region of F. ARM residues that are either involved in hydrogen bonds or salt bridges with BRCA2, or buried by more than 30 Å^2^ at the interface with BRCA2, are represented by colored sticks and labeled. Residues identified in only one of these two similar interfaces are labeled in italics.

**Supplementary Figure S8.**
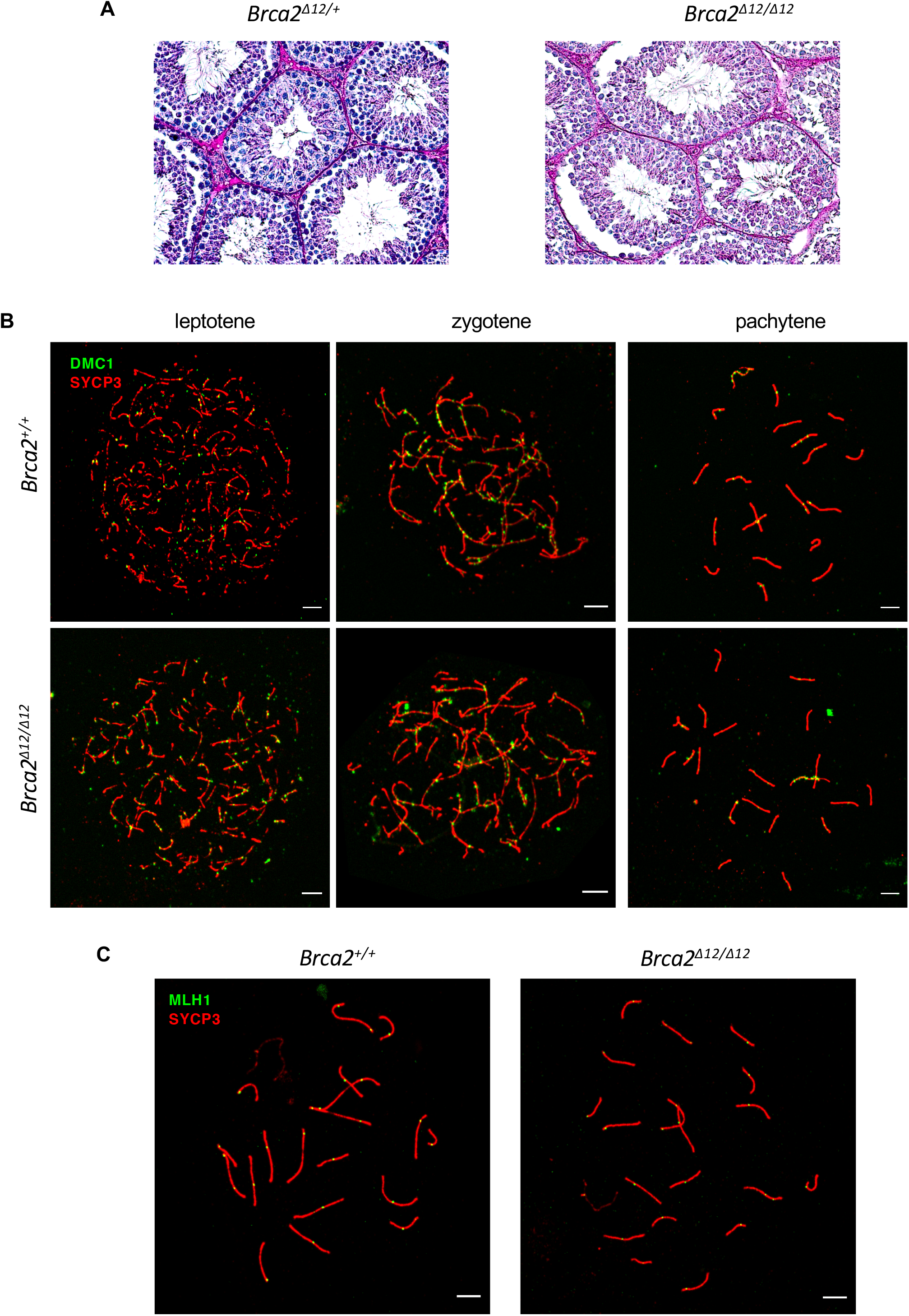
**(A)** Representative histological images of testis cross-sections. (**C**) Representative images used for DMC1 foci quantification. (**C**) Representative images used for MLH1 foci quantification.

**Supplementary Figure S9.**
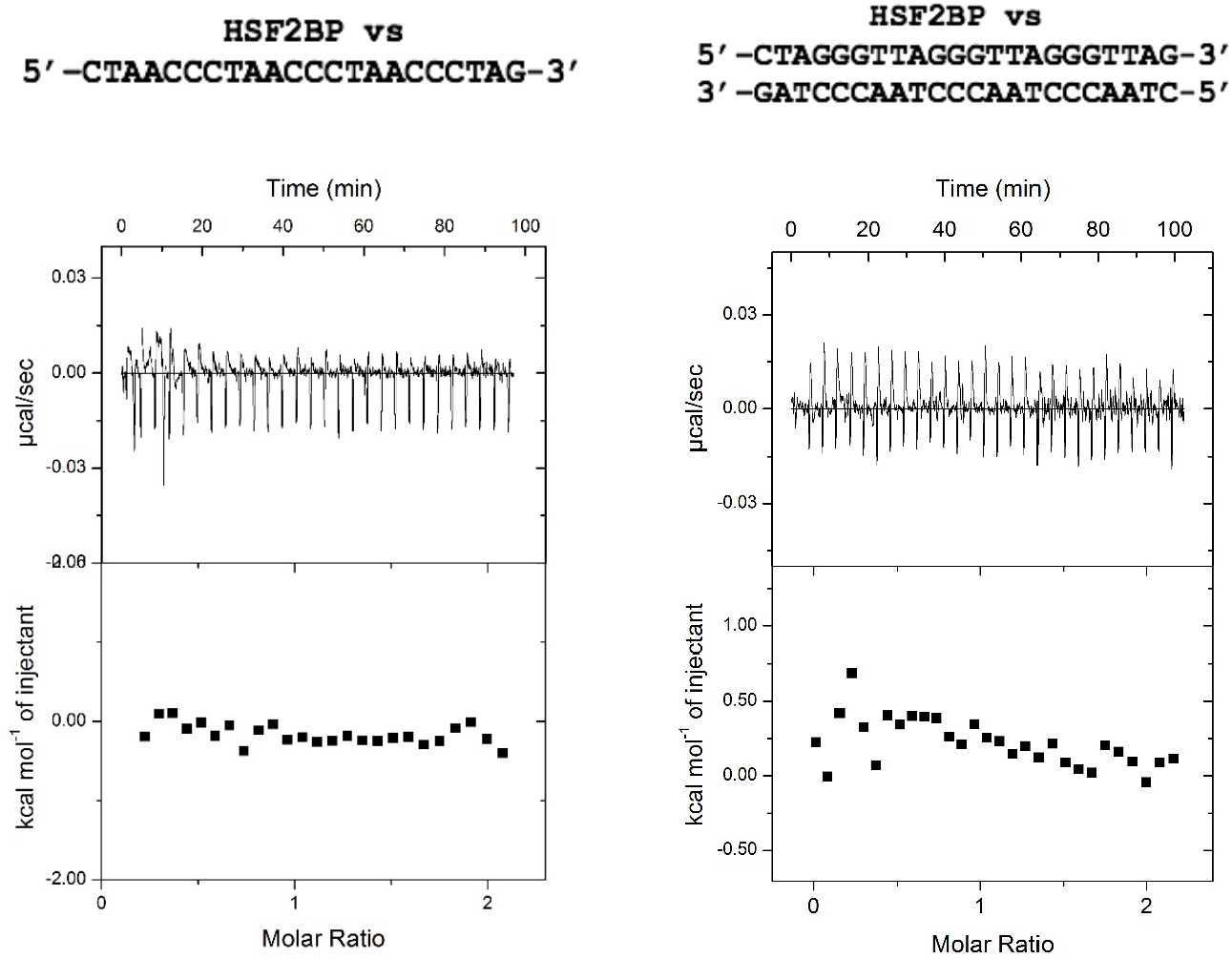
ITC analysis of the interactions between HSF2BP and DNA. HSF2BP (7-9 μM in the instrument cell) did not interact with ssDNA and dsDNA oligonucleotides (70-90 μM in the instrument syringe).

Supplementary Movies 1-3. Live cell recording of HSF2BP-GFP diffusion in the nuclei of mouse ES cells with the following genotypes: (1) *Brca2^+/+^Hsf2bp^GFP/GFP^;* (2) *Brca2^Δ12/Δ12^Hsf2bp^GFP/GFP^;* (3) *Brca2^Δ12-14/Δ12-14^Hsf2bp^GFP/GFP^*.

